# Genome diversity and phylogeny of the section *Alatae* of genus *Lemna* (Lemnaceae), comprising the presumed species *Lemna aequinoctialis, Le. perpusilla* and *Le. aoukikusa*

**DOI:** 10.1101/2025.01.17.632899

**Authors:** Anton Stepanenko, Luca Braglia, Jörg Fuchs, Veit Schubert, Phuong TN Hoang, Yuri Lee, Guimin Chen, Silvia Gianì, Laura Morello, Ingo Schubert

## Abstract

- The Section *Alatae* of genus *Lemna* of the monocotyledonous aquatic duckweed family (Lemnaceae) consists of rather diverse accessions with unknown phylogeny and unclear taxonomic assignment. In contrast to other duckweeds, some *Alatae* accessions, in addition to mainly vegetative propagation, produce readily flowers and viable seeds. We analyzed the genomic diversity and phylogenetic relationship of 52 *Alatae* accessions.
- For this purpose, we applied multiple molecular and cytogenetic approaches, including plastid and nuclear sequence polymorphisms, chromosome counting, genome size determination and genomic *in situ* hybridization in combination with geographic distribution.
- We uncovered ploidy variation, recurrent hybridization, and backcrosses between diploid species and their hybrids. The latter successfully spread over three continents.
- The results elucidate the evolution of *Alatae* accessions (summarized in Fig. 7) and explain the difficult taxonomic assignment of distinct accessions. Our study might be an example for analogous studies to resolve the hitherto unclear relationships among accessions of the duckweed genera *Wolffiella* and *Wolffia*.

## Introduction

The aquatic monocotyledonous duckweeds, the family Lemnaceae, Martinov, comprise five genera with so far 35 species and two hybrid species (Appenroth et al., 2024; Bog et al., 2019). Despite their small size and reduced morphology, duckweeds are remarkably diverse and their genetic, physiological and biochemical features, their development, evolution and practical applications are of increasing interest (for review see Cao et al., 2018; Fourounjian et al., 2020; Acosta et al., 2021). While the phylogeny of the genus *Spirodela* Schleid. with two species, and the monospecific genus *Landoltia* Les & Crawford is obvious, this is not the case for accessions of the remaining genera *Lemna* L.*, Wolffiella* Hegelm. and *Wolffia* Horkel ex Schleid. Despite the mainly vegetative propagation of duckweeds, within the genus *Lemna* ploidy variants and so far two hybrid species, *Lemna japonica* Landolt (*Le*. *minor* L. *× Le*. *turionifera* Landolt, see Braglia et al., 2021a; 2021b; Ernst et al., 2023, preprint) and *Le*. *mediterranea* Braglia et Morello (*Le*. *minor × Le*. *gibba* L. and reciprocal, see Braglia et al., 2024) have been proven.

Among the genus *Lemna, Le. aequinoctialis* Welw. (lesser duckweed) has great prospects for practical use as feed for fish, poultry, ruminants and pigs, as food for humans, as a bioreactor for proteinaceous compounds (Ma et al., 2018; Nati et al., 2024) as well as for bioethanol production (Faizal et al., 2021) and for wastewater remediation (Zhou et al., 2018; Hu et al., 2019; Shi et al., 2020; Cai et al., 2022; Toyama et al., 2024). This is due to its wide geographical distribution in tropical and subtropical climates (Landolt, 1980), and nowadays even in the temperate zones of Europa (Vélez-Gavilán, 2022; Fedoniuk et al., 2022), its rapid biomass generation, valuable composition, and lack of cytotoxic effects (Nati et al., 2024). The potential for practical use of the sister species *Le. perpusilla* Torr. (minute duckweed) is more scarce due to the small number of available accessions in the public collections and to its limited distribution (in the Middle and Eastern states of the USA and Canada (Landolt, 1986; Tippery & Les, 2020). Some researchers claimed its occurrence also for India, China, Korea and Saudi Arabia (Halder and Venu, 2012; Xu et al., 2018; Al-Dakhil et al., 2021; Lee et al., 2022). However, the approaches used to identify *Le. perpusilla* outside North America are questionable. Thus, there are only a few reports on its use for phytoremediation (Clark et al., 1981; Tang et al., 2013).

*Le. aequinoctialis* and *Le. perpusilla* are the only currently recognised species in the section *Alatae*, with relatively frequent fruiting. They differ morphologically regarding the number of indistinct ribs of seeds and the seed behavior after ripening (*Le. perpusilla* seeds stay within the fruit wall after ripening, while *Le*. *aequinoctialis* seeds fall out) (Kandeler & Hügel, 1974; Landolt, 1986). In earlier studies, *Le. aequinoctialis* and *Le. perpusilla* were considered synonymous (Daubs, 1965; Den Hartog & Van der Plas, 1970), but Landolt (1980) recognised them as separate species, as has been confirmed by allozyme studies (Crawford et al., 2001), flavonoid and anatomical-morphological data (Les et al., 1997), and AFLP (Bog et al., 2010). In addition, some *Alatae* accessions were referred to either *Le. aequinoctialis* or *Le. perpusilla* in different publications, and occasionally, the old name *Le*. *paucicostata* Hegelm., then synonymized to *Le. aequinoctialis* (Landolt, 1986; Sree et al., 2016), is used for *Alatae* accessions, which caused confusion regarding the identification of these species.

In addition, some researchers distinguish a third species of the *Alatae* section, *Le. aoukikusa* Beppu et Murata, which has been found in Japan. Unlike *Le. aequinoctialis*, *Le. aoukikusa* can survive in colder climates and displays some morphological differences regarding frond thickness, root cap, anther size, flower development and self-compatibility (Beppu & Takimoto, 1981; Beppu et al., 1985; Lee et al., 2024). However, it is currently not clear whether the corresponding accessions represent a separate species, a subspecies or simply a geographically isolated *Le. aequinoctialis* population with variable morphological characteristics.

Therefore, deeper studies of the biodiversity and phylogeny of the *Alatae* section are required. Most of the previous phylogenetic studies of Lemnaceae included only a small number of presumed *Le. aequinoctialis* and *Le. perpusilla* accessions (Bog et al., 2010; Chen et al., 2022; Tippery et al., 2015; Braglia et al., 2021b), or only specimens collected from geographically restricted areas (Wang et al., 2010; Tang et al., 2014; Borisjuk et al., 2015; Tang et al., 2015; Xu et al., 2018; Barbosa Neto et al., 2019; Lee et al., 2024). The obtained data were not sufficient for a comprehensive assessment of the diversity of these species, the more so as *Le. aequinoctialis* accessions are morphologically rather variable (Barbosa-Neto et al., 2019; Lee et al., 2024).

Selecting suitable methods to study diversity and phylogenetic relationship of duckweeds is challenging. Chloroplast barcoding is a relatively inexpensive and easy-to-implement method (Wang et al., 2010; Borisjuk et al., 2015). Therefore, it is used in most phylogenetic studies on duckweeds (Tang et al., 2014; Tang et al., 2015; Xu et al., 2018; Barbosa-Neto et al., 2019; Braglia et al., 2021a, 2024; Chen et al., 2022; Lee et al., 2024). One of its disadvantages is the inability to identify interspecific hybrids. Another one is the low level of polymorphism between closely related accessions. The latter problem can be overcome by applying multiple marker sequences (Chen et al., 2022). Using nuclear markers is another approach to study the biodiversity of duckweeds. For this purpose, 35S rDNA ITS1 and ITS2 (Tippery et al., 2015), 5S rDNA NTS (Chen et al., 2024), AFLP (Bog et al., 2010), SSR (Xu et al., 2018), Tubulin-Based Polymorphism (TBP; Braglia et al., 2021a, b, 2024) and genotyping-by-sequencing have been applied (Bog et al., 2020c).

Cytogenetic parameters, such as chromosome number and genome size, have proven useful for duckweed taxonomy, highlighting the presence of different cytotypes within the same species (Landolt, 1980; Hoang et al., 2019; Bog et al., 2020b). In addition, genomic *in situ* hybridization (GISH) confirmed interspecific hybrids (Ernst et al., 2023, preprint). All of these approaches have advantages and disadvantages and no single one alone is of sufficiently informative value (Bog et al., 2022).

Under these circumstances, this study aimed to resolve the phylogenetic relationship of presumed *Le. aequinoctialis, Le. perpusilla* and *Le. aoukikusa* accessions by several independent approaches, including plastid and nuclear markers, genome size, chromosome counts and genomic *in situ* hybridization, applied to about 50 accessions from different continents and climatic zones.

## Materials and Methods

### Plant material

We included in our study 52 *Alatae* accessions from different continents (Europe, Asia, America and Africa) and two *Le. tenera* clones. The full list of accessions and research conducted on them is given in Supporting Information Table S1. The analyzed *Le. aequinoctialis, Le. perpusilla* and *Le. aoukikusa* accessions belong to author’s collections or were kindly provided by Klaus J. Appenroth, Nikolai Borisjuk, Manuela Bog, Sowjanya Sree and Eric Lam. The accessions were sterilized as described (Appenroth, 2023). The sterile clones were maintained on solid N medium (Appenroth, 1996) with 0.6% of Gelrite. To obtain plant material for DNA extraction and metaphase chromosome preparation, duckweeds were grown on liquid SH medium (Schenk & Hildebrandt, 1972) with 5 gL^-1^ sucrose. For some accessions which meanwhile were lost from collections (EL031, NBFu94, 8656), DNA preparations were preserved and included in this study for analyses of plastid and ITS polymorphisms.

### DNA extraction and chloroplast barcoding

Total DNA was extracted from plant tissue using a modified CTAB method (Murray & Thompson, 1980). The PCR amplification for barcoding was carried out as recommended by the CBOL Plant Working Group (2009), described in Wang et al. (2010), using specific primers for the *atpF*–*atpH* and *psbK*–*psbI* barcode. Following amplification, the DNA fragments were purified using the NucleoSpin Gel and PCR Clean-up Kit (Macherey-Nagel, Germany) and sequenced using a custom service provided by LGS Genomics GmbH (Berlin, Germany). The obtained forward and reversed sequences were assembled and analyzed using the CLC Main Workbench (Version 6.9.2, Qiagen) software. The obtained sequences were deposited in GeneBank (for accession numbers see Supporting Information Table S1).

### Cloning and sequencing of ribosomal RNA genes

For analysis of the ITS1–5.8S–ITS2 region of *Alatae* accessions, the specific DNA fragments were amplified by PCR from the same samples of total DNA as for barcoding. In the standard PCR, we used two primers (18S–F1 and 25S–R1) specific for 18S and 25S rDNA (Supporting InformationTable S2), respectively. The protocol was optimized to amplify GC-rich regions (Stepanenko et al., 2022). After cutting out the gel with PCR products and DNA purification using NucleoSpin Gel and PCR Clean-up Kit (Macherey-Nagel, Germany), the generated fragments were cloned into the vector pMD19 (Takara, Dalian, China). After collecting the positive bacterial clones, plasmids were extracted using NucleoSpin Plasmid Kit (Macherey-Nagel, Germany), and were sequenced using a custom service provided by LGS Genomics GmbH (Berlin, Germany).

The obtained sequences were assembled and analyzed using the CLC Main Workbench (Version 6.9.2, Qiagen) software. At least three ITS1 and ITS2 sequences along with the 5.8 rDNA sequence were obtained for each duckweed clone to produce a consensus sequence. The obtained consensus sequences were deposited in GeneBank (for accession numbers see Supporting Information Table S1).

### Phylogenetic Analysis

The maximum-likelihood phylogenetic trees for phylogenetic analysis of ITS and plastid sequences were constructed applying the NGPhylogeny web-service (https://ngphylogeny.fr) (Lemoine et al., 2019) using MAFFT Multiple Sequence Alignment (Katoh & Standley, 2013) and PhyML algorithm with Smart Model Selection (Guindon et al., 2010). Cleaning aligned sequences was made by utilizing BMGE tools (Criscuolo & Gribaldo, 2010). Bootstrap support (BS) (Felsenstein’s bootstrap proportions (FBPs) + transfer bootstrap expectation (TBE)) was estimated with 100 boot-strap replicates. iTOL (https://itol.embl.de) and used for displaying and annotating the generated phylogenetic trees (Letunic & Bork, 2021).

### Genome size measurement

For flow cytometric genome size measurements fresh leafy fronds were chopped together with an appropriate amount of leaf tissue of one of the reference standards (*Raphanus sativus* cultivar ‘Voran’, Gatersleben Genebank accession number: RA 34 [1.11 pg/2C]; *Lycopersicon esculentum* Mill. convar. *infiniens* Lehm. var. *flammatum* Lehm., cultivar ‘Stupicke Rane’, Gatersleben Genebank accession number: LYC 418 [1.96 pg/2C] or *Glycine max* (L.) Merr. convar. *max* var. *max*, cultivar ‘Cina 5202’, Gatersleben Genebank accession number: SOJA 392 [2.21 pg/2C]) with a sharp razor blade and using the ‘CyStain PI Absolute P’ nuclei extraction and staining kit (Partec-Sysmex) according to the manufacturer’s instruction. After filtering the suspension through a 50 µm mesh (CellTrics, Sysmex-Partec) samples were analyzed on a CyFlow Space flow cytometer (Partec-Sysmex, Görlitz, Germany). The DNA content (pg/2C) was calculated based on the values of the G1 peak means of sample and reference species, and the corresponding genome size (Mbp/1C) according to Dolezel et al. (2003).

### Mitotic chromosome preparation

The plants were grown in SH medium (Schenk & Hildebrandt, 1972) with 5 g<u>L</u>^-1^ sucrose until new daughter fronds had developed. The fronds were collected, treated in 0.02 M 8-hydroxyquinoline at 37°C for 1 h and then fixed in fresh 3:1 (absolute ethanol: acetic acid) for 24 h. The fixed samples were washed twice in 10 mM Na-citrate buffer pH 4.6 for 10 min each, before and after softening in 2 ml PC enzyme mixture [2% pectinase and 2% cellulase in Na-citrate buffer] for 60 min at 37°C, before maceration and squashing in 60% acetic acid. After freezing in liquid nitrogen, the slides were treated with pepsin, (50 µgml^-1^ pepsin in 0.01N HCl, 5 min at 37°C), post-fixed in 4% formaldehyde in 2x SSC for 10 min, rinsed twice in 2x SSC, 5 min each, dehydrated in an ethanol series (70, 90 and 96%, 2 min each) and air-dried.

### Probe labelling

The DNA probes specific for the genomes of diploid *Le. aequinoctialis* and *Le. perpusilla,* respectively, were prepared following the procedure described by Hoang & Schubert (2017). First, genomic DNA isolated from *Le. aequinoctialis* (2018) and *Le. perpusilla* (Bog0007) was sonicated to obtain fragments of 500–1000 bp. This DNA was used as template for labelling with specific fluorescent dyes. The DNA was subjected to nick-translation using the Alexa594 NT Labeling Kit and the Atto488 NT Labeling Kit (Jena Bioscience, Jena, Germany). After precipitation with 96% ethanol, probe pellets were dissolved in 100 μl hybridization buffer (50% (v/v) formamide, 20% (w/v) dextran sulfate in 2 × SSC, pH 7).

### Genomic in situ hybridization (GISH)

GISH with genomic DNA of *Le. aequinoctialis* and *Le. perpusilla* was applied to well-spread chromosome preparations of the species of probe origin and those of presumed hybrids. Probes were denatured at 95°C for 5 min and chilled on ice for 10 min before adding 10 µl of each probe per slide. Then, the mitotic chromosome preparations were denatured together with the probes on a heating plate at 80°C for 3 min, followed by incubation in a moist chamber at 37°C for at least 16 h. Post-hybridization washing and signal detection were done as described (Lysak et al., 2006) with minor modifications. Widefield fluorescence microscopy for signal detection followed Cao et al. (2016).

### Super-resolution microscopy

To analyze the ultrastructure of chromatin beyond the classical lateral Abbe-Rayleigh limit of ∼250 nm, spatial structured illumination microscopy (3D-SIM) was performed to achieve a lateral resolution of ∼120 nm (super-resolution, attained with a 488 nm laser). We used an Elyra 7 microscope system equipped with a 63×/1.4 Plan-Apochromat objective and the ZENBlack software (Carl Zeiss GmbH). Image stacks were captured separately for each fluorochrome using 405 nm (DAPI), 488 nm (Atto488) and 561 nm (Alexa594) laser lines for excitation and appropriate emission filters (Kubalova et al., 2021; Weisshart et al., 2016). Zoom-in sections are presented as single slices to detect the subnuclear chromatin structures at the super-resolution level. Wide-field images were processed based on the SIM raw data. Chromosome counting was done within the spatial image stacks.

### TBP profiling

The capillary electrophoresis Tubulin Based Polymorphism (CE-TBP) method was carried out according to the procedure defined by Braglia et al. (2023). The 1^st^ and 2^nd^ intron amplifications from 30 ng of genomic DNA were performed twice per clone. Control PCR reactions, without DNA template, were always included in any experiment. The signal intensity of the PCR amplicons was checked on a 2% agarose gel and the diluted samples were prepared for the capillary electrophoresis separation according to Braglia et al. (2016). Data analysis and collection were performed using the Gene Mapper Software v.5.0 tools (Thermo Fisher Scientific, Inc., Waltham, MA, USA) according to the parameters defined by Braglia et al. (2023). The size (in base pairs) and the height (in relative fluorescence units = RFUs) of each peak composing the CE-TBP pherogram of each clone were collected into Microsoft Office Excel files. The numerical data sorted according to the peak size was considered comparing the CE-TBP profiles of the analyzed samples. A presence/absence matrix (1/0 respectively) was obtained by the peak scoring of both intron regions (1^st^ and 2^nd^). The Jaccard’s index genetic similarities among clones were estimated, and a neighbor-joining (NJ) dendrogram, with 1000 replicates (bootstrap analysis), was computed using the open-source software package Past v.4.14 (Hammer et al., 2001).

## Results

### Genotyping by chloroplast barcoding

We genotyped 52 *Alatae* and two *Le. tenera* accessions by direct sequencing of the chloroplast gene spacer sequences of *atpF*–*atpH* and *psbK*–*psbI* (Supporting Information Figs. S1 and S2). Since the analyses of single spacers did not give unambiguous results, we combined *ATP* and *PSB* analyses using *Le. tenera* Kurz as an outgroup species. To confirm the identity of *Le. tenera*, we sequenced the *atpF*–*atpH* spacers for accessions 9020 and 9024. *psbK*–*psbI* spacers were obtained from NCBI (accession numbers KJ136049 and KJ136048, respectively) for the same *Le. tenera* accessions. The combined data matrix for the presumed species *Le. aequinoctialis*, *Le. perpusilla* and *Le. aoukikusa* contained 972 characters, divided into two partitions: 1–465 for *atpF*–*atpH* and 466–972 for *psbK*–*psbI*, of which 892 were constant, 17 variable characters were parsimony uninformative and 63 were parsimony informative. The combined data matrix for all tested accessions including *Le. tenera* as outgroup contained 847 constant, 21 variable but parsimony uninformative, and 104 parsimony informative characters. The resulting phylogenetic tree is shown in the left part of Fig. **1**.

**Figure 1.**
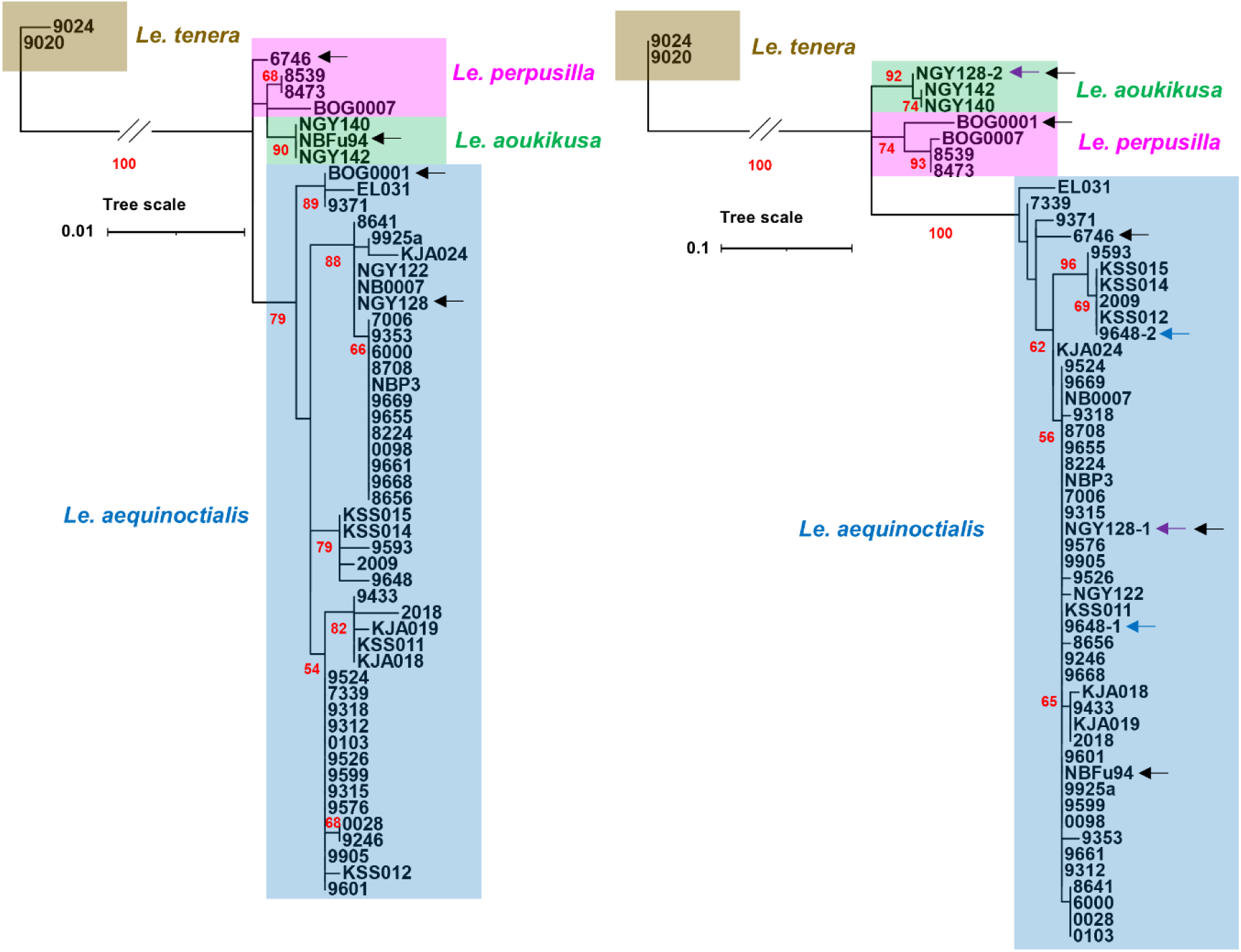
Phylogeny of presumed *Le. aequinoctialis, Le. perpusilla, Le. aoukikusa* and *Le. tenera* accessions based on *atpF*–*atpH* and *psbK*–*psbI* intergenic spacer sequences **(left)** and on ITS1–5.8rDNA–ITS2 sequences **(right)**. Black arrows indicate accessions with different positions in barcoding and ITS trees; colored arrows indicate accessions for which different ITS sequences were found. Bootstrap values >50 are shown by red numbers at the branches.

*Le. aequinoctialis* clones form a distinct cluster with several subclusters. Notably, one of the subclusters contains BOG0001, which previously was assigned to *Le. perpusilla* based on tubulin gene polymorphism (Braglia et al., 2021b). In addition to this accession from the South-Eastern United States, the subcluster contains 9371 and EL031 from Venezuela and Brazil, respectively (for geographic origin of accessions see also Fig. **6** and Supporting Information Table S1).

The remaining accessions form one cluster with strongly bootstrap-supported branches. One branch contains two presumed *Le. aoukikusa* accessions from East Asia (NGY140, NGY142) and NBFu94, originally classified as *Le. aequinoctialis*; another one, three *Le. perpusilla* accessions from the Eastern USA of which BOG0007 differs from the other two. The presumed *Le. aequinoctialis* accession 6746 from the western part of the USA forms a separate branch. This clone shares five specific SNPs/indels with *Le. perpusilla* and only two SNPs with *Le. aequinoctialis*.

### Genotyping by ITS1 and ITS2

The variable spacer sequences between the 18S and 5.8S rDNA (ITS1) and between 5.8S rDNA and 25S rDNA (ITS2) are useful nuclear markers for phylogenetic studies of close relatives. To amplify these sequences, we developed a set of primers localized on 18S and 25S rDNA. The ITS1–5.8S–ITS2 regions were cloned and sequenced for 52 *Alatae*, and two *Le. tenera* accessions as outgroup (Supporting Information Fig. S3).

The combined data matrix for *Le. aequinoctialis, Le. perpusilla* and *Le. aoukikusa* included 724 characters divided into three partitions: 1–307 for ITS1, 308–465 for 5.8 rRNA gene and 466– 724 of which 662 were constant, 19 were variable but parsimony uninformative, and 43 were parsimony informative. The combined data matrix for all accessions, including *Le. tenera* as an outgroup, contains 589 constant characters, 15 variable but parsimony uninformative, and 120 parsimony informative characters. The resulting phylogenetic tree is shown in the right part of Fig. **1**.

Bioinformatic and phylogenetic analyses revealed compelling differences between the ITSs of *Le. aequinoctialis*, *Le. perpusilla, Le. aoukikusa* and *Le. tenera* accessions. The tested accessions displayed only one type of ITS1 and ITS2, except for 9648 and NGY128, which showed two types of ITS sequences, suggesting their hybrid nature (9648 intraspecific between different *Le. aequinoctialis* accessions, and NGY128 between *Le. aequinoctialis* and *Le. aoukikusa*).

Apart from the out-group accessions of *Le. tenera,* the two *Le. aoukikusa* accessions and NGY128 formed a separate cluster, distinct from those of *Le. aequinoctialis* and *Le. perpusilla*, but closer to *Le. perpusilla*. It should be noted that NBFu94 from China, assigned to *Le. aoukikusa* according to chloroplast barcoding results, belongs to *Le. aequinoctialis*, but its ITS marks it as another hybrid candidate, a potential backcross of *Le. aoukikusa* with *Le. aequinoctialis* as paternal ancestor. Because this clone is no longer available, no further investigation was possible. Remarkably, also the presumed *Le. aequinoctialis* accession 6746 and the presumed *Le. perpusilla* accession BOG0001 showed discordant results by chloroplast barcoding and ITS genotyping. Thus, they are also hybrid candidates. That sometimes ITS sequences are not detected from both presumed parental ancestors could be due to frequent uniparental loss of one ancestor’s rDNA.

Three *Le. perpusilla* accessions and BOG0001 (assigned to *Le. aequinoctialis* by barcoding) formed a distinct cluster, with BOG0001 branching apart from the other three.

The presumed *Le. aequinoctialis* accessions form a separate, rather homogeneous cluster with strong support. Notably, three American clones, 6746, 9371 and EL031, belong to this cluster but are quite variable and different from the main group of *Le. aequinoctialis*.

To further study discrepancies between barcoding and ITS trees as well as to confirm potential hybrids, we measured genome sizes, counted chromosomes and performed GISH experiments.

### Genome size measurements

Genome size was measured by flow cytometry in at least three replicates for each tested accession. According to their genome size, the presumed *Le. aequinoctialis* accessions are divided into distinct groups (Fig. **2**, Supporting Information Table S1). The first largest group contains 24 clones with a genome size ranging from 446 to 510 Mbp, except for accession 9371 with only 413 Mbp. The second group consists of potentially triploids, displaying a genome size variation of 14%. To the accessions of triploid genome sizes belongs 9648, suspected to be an intraspecific hybrid, based on two ITS variants. The three diploid *Le. perpusilla* accessions (8473, 8539 and BOG0007) have deviating genome sizes of 520, 522 and 585 Mbp, respectively. The five potentially tetraploid clones vary also regarding their genome size, ranging from 890 Mbp for 6746 to 1018 Mbp for BOG0001, both of which are potential hybrids according to discordant results in chloroplast barcoding and ITS genotyping. The remaining three tetraploid accessions (NGY142, NGY140 and NGY128) were previously assigned to *Le. aoukikusa* or to *Le. aequinoctialis* (NGY128), respectively (Lee et al., 2024).

**Figure 2.**
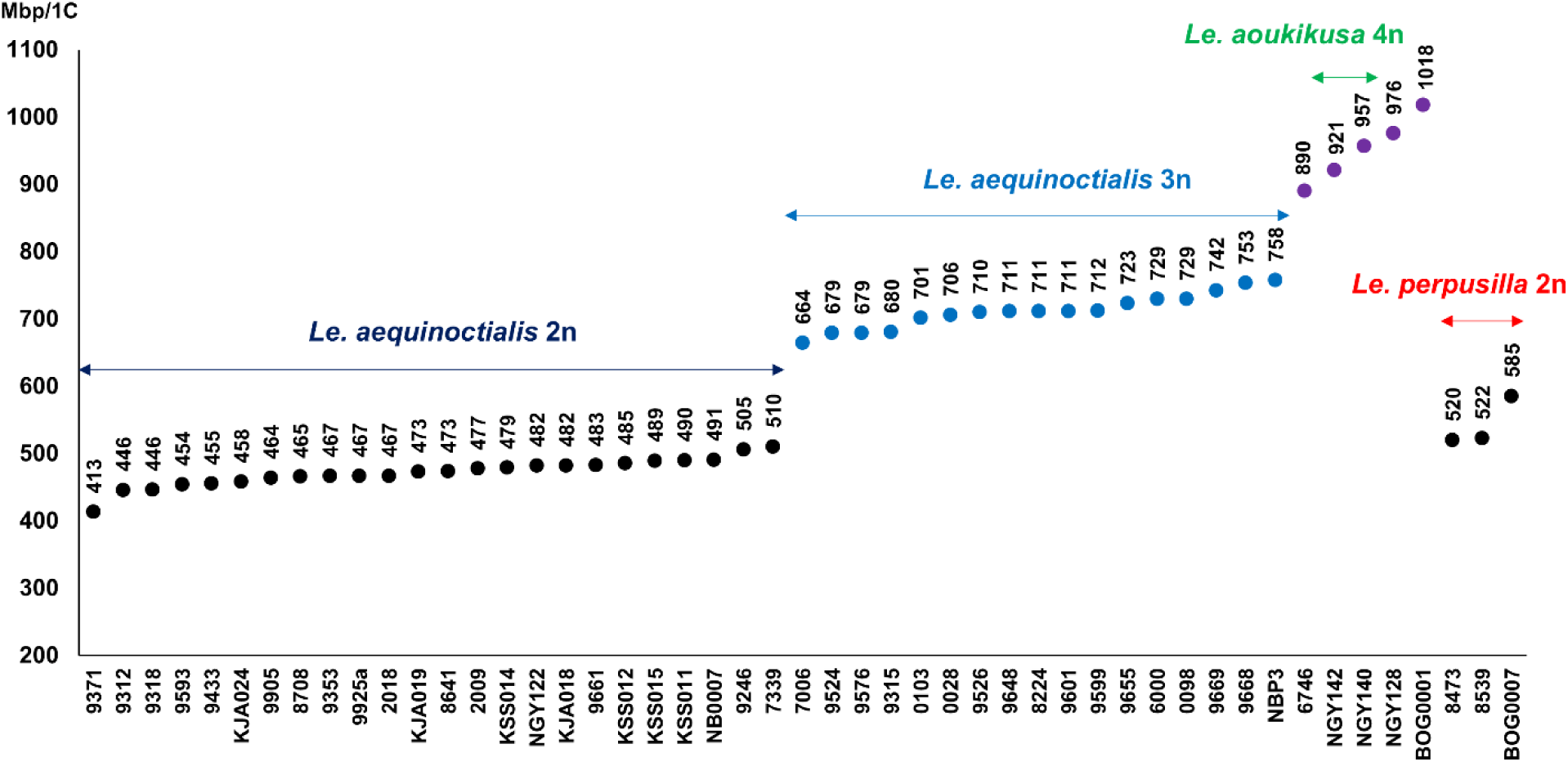
Genome size measuring revealed potentially diploid (black), triploid (blue) and tetraploid (violet) *Alatae* accessions. Species assignment is according to what has been confirmed in our study.

To confirm the ploidy variants, some accessions were selected for chromosome counting. To test for potential allopolyploidy, tri- and tetraploid accessions were submitted to GISH.

### Chromosome counting

We selected for chromosome counting diploid accessions of *Le. aequinoctialis* (NB0007, 9925a) and *Le. perpusilla* (BOG0007, 8473), triploid (9648, 9668, 7006, 9526) and tetraploid (BOG0001, NGY142, 6746, NGY128) *Alatae* accessions from different groups based on molecular marker diversity. The number of chromosomes was counted on at least three metaphases for each selected accession.

The diploid accessions displayed invariably 42 predominantly small chromosomes (2n=42) after DAPI staining. These results are consistent with previous counts for the *Le. aequinoctialis* accession 2018 (Hoang et al., 2019). Sixty-three chromosomes were counted for all tested triploid accessions and 84 for tetraploid ones (Fig. **3**). These data reflect previously reported variability of chromosome number for *Le. aequinoctialis* (for review see Hoang et al., 2022). Intermediate numbers, such as 50, 60, 70 and 80 chromosomes as previously reported for some *Le. aequinoctialis* accessions (Urbanska, 1980) are not supported. Due to their small size and often lacking structural details, such as primary (centromere) or secondary (nucleolus-organising region [NOR]) constrictions, these chromosomes were barely individually distinguishable and it remains an open question whether they are mono- or holocentric.

**Figure 3.**
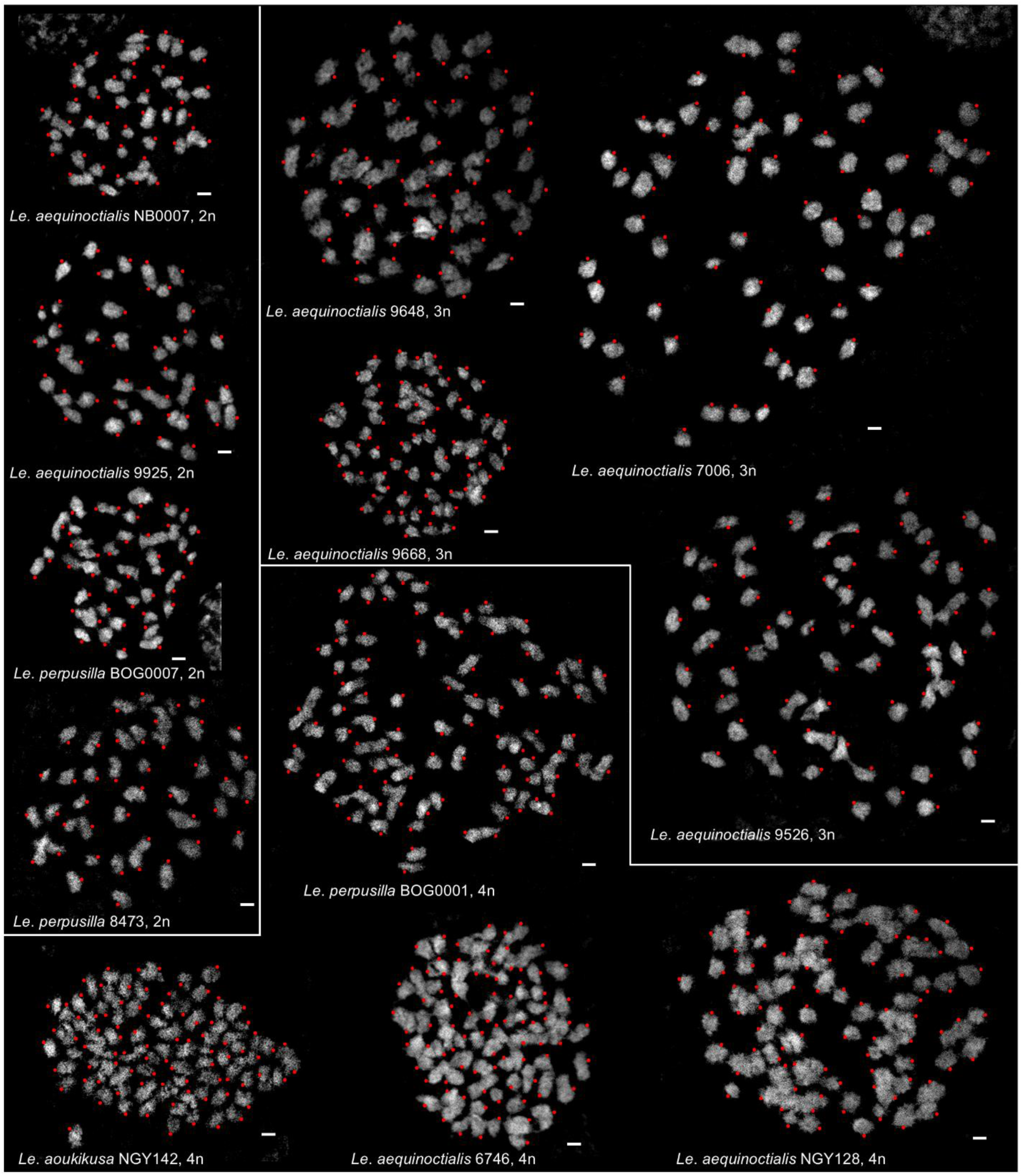
Chromosome spreads of selected diploid (2n=42), triploid (2n=63) and tetraploid (2n=84) *Alatae* accessions.

### Genomic in situ hybridization (GISH)

For the analysis of potential allopolyploidy, GISH with genomic DNA of the potential parental performed with differently labelled genomic DNA from two diploid accessions, *Le. aequinoctialis* 2018 and *Le. perpusilla* BOG0007, against the chromosomes of both accessions (Fig. 4). This resulted, as expected, in specific labelling of the chromosomes by the DNA of the corresponding accession with a low level of cross-hybridization, possibly indicating their close relatedness (see zoomed inserts in Fig. **4**).

**Figure 4.**
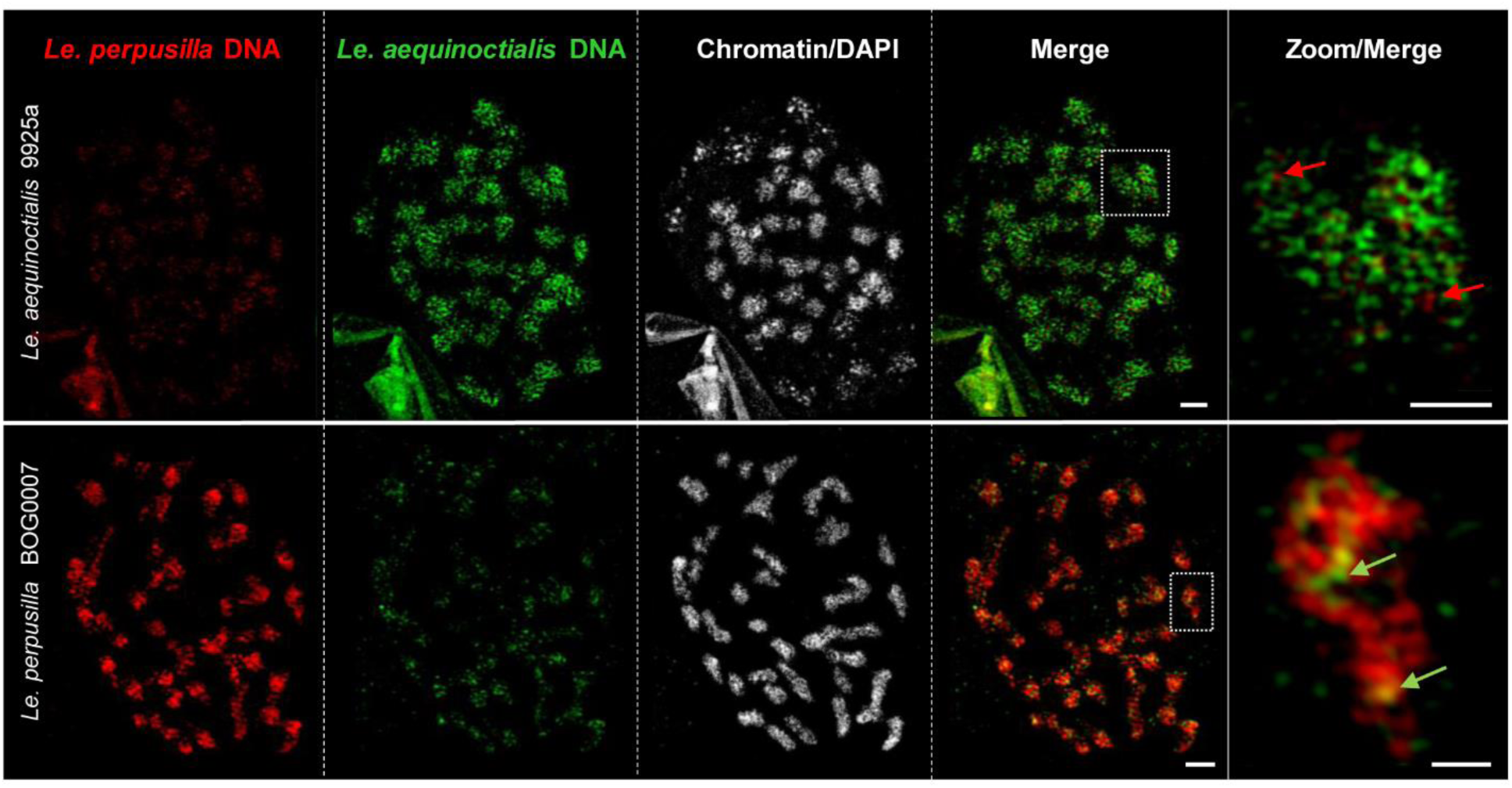
Fluorescence *in situ* hybridization with genomic DNA of *Le. perpusilla* (red) and *Le. aequinoctialis* (green) on mitotic chromosomes of diploid *Le. aequinoctialis* 9925a and *Le. perpusilla* BOG0007. The enlarged insets show cases of cross-hybridization (arrows). Bars in complete metaphase cells: 2 µm; in enlarged regions: 0.5 µm.

GISH was then performed on triploid (9669, 7006) and tetraploid accessions (NGY142, NGY128, BOG0001, 6746), the latter three suspected to be hybrids based on discordant barcoding and ITS data (Fig. **1**). All of them were confirmed as interspecific hybrids between *Le. aequinoctialis* and *Le. perpusilla* (Fig. **5**), including NGY142, which was described previously as fertile *Le. aoukikusa* (Lee et al., 2024).

**Figure 5.**
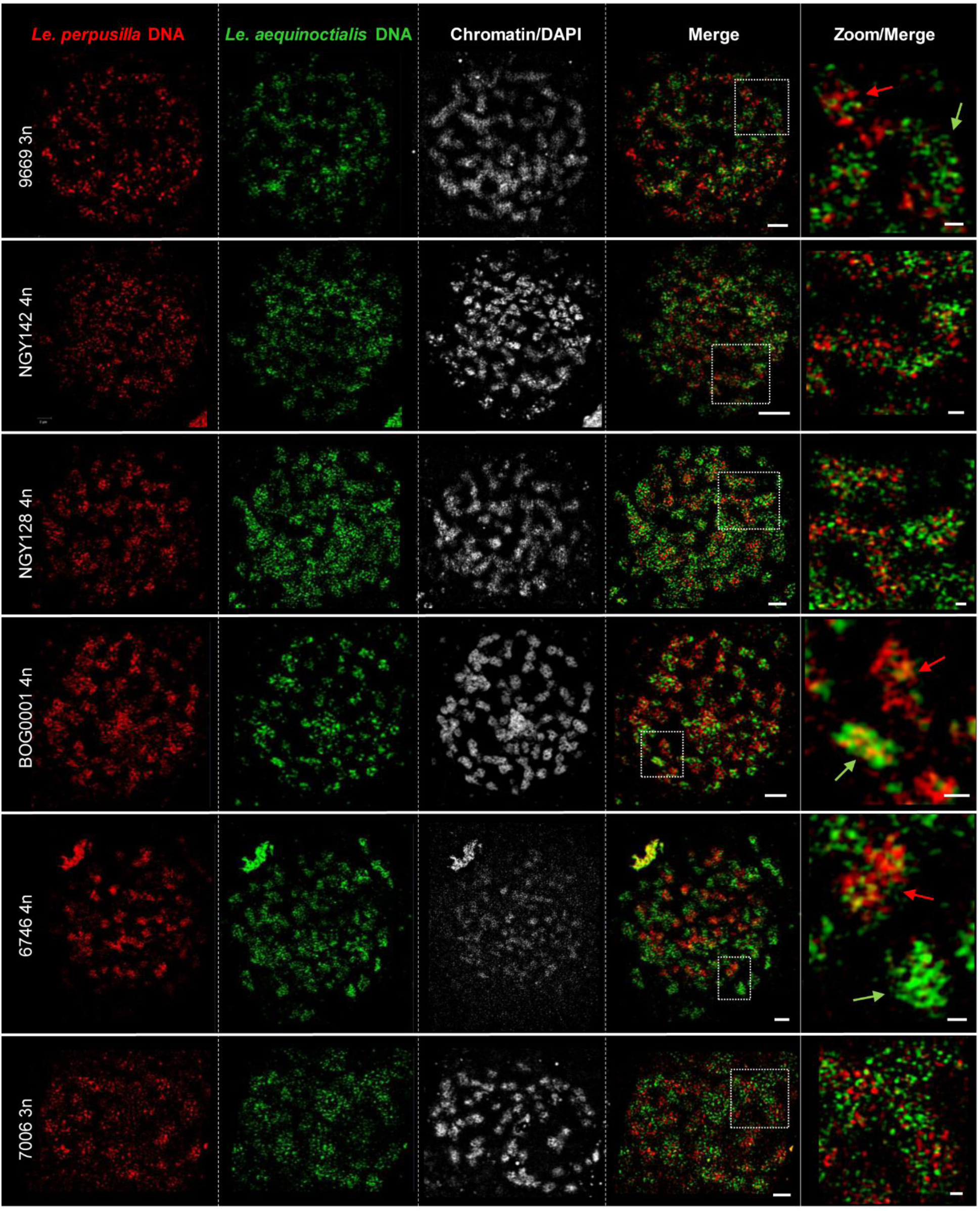
Mitotic metaphases of tri- and tetraploid accessions after GISH using genomic *Le. perpusilla* (red) and *Le. aequinoctialis* (green) DNA and applying SIM. The intensity of red and green signals counted on SIM images was close to a 1:1 ratio for exemplarily counted accessions NGY142, NGY128 and 6746. The enlarged regions show cross-hybridizations (arrows) in enlarged regions: 0.5 µm.

Cross-hybridization, that means red signals on predominantly green chromosomes and *vice versa,* suggests somatic recombination between homeologs during double-strand break (DSB) repair via gene conversion as evidenced between sister chromatids of the field bean (Schubert et al., 2011). The *in situ* cross-hybridization frequency likely increases with the age of the hybrid.

### Genotyping by tubulin-based polymorphisms (TBP)

The combined data for genome size, chromosome number (=ploidy level), GISH and geographic origin are integrated into the phylogenetic tree derived from TBP analyses for the *Alatae* accessions (Fig. **6**). At its first node, the TBP tree clearly separates three diploid *Le. perpusilla s*.*s*. accessions and relatively young (compared to *Le. aoukikusa*) tri-(7006) and tetraploid (BOG0001 and 6746) hybrids with *Le. aequinoctialis* (as confirmed by GISH), from all other clones (pink dot). Accessions 7006 and BOG0001 have *Le. aequinoctialis,* and 6746 has *Le. perpusilla*, as maternal parent, as suggested by chloroplast barcoding.

A second node (green dot) separates *Le. aequinoctialis s*.*s*. from its ancient tetraploid hybrids with *Le. perpusilla*, *Le. aoukikusa* (NGY140, NGY142), and the tri- and tetraploid backcross accessions with *Le. aequinoctialis* as maternal parent, according to barcoding results (Fig. **1**). The allotriploids are supposed to have received the *Le. perpusilla* genome through hybridization of *Le*. *aoukikusa* with diploid *Le. aequinoctialis*, as suggested by their closer proximity to *Le. aoukikusa.* than to *Le*. *perpusilla.* For the tetraploid accession NGY128, hybridity between *Le. aequinoctialis* (as plastid donor) and *Le. aoukikusa,* is supported by the heterozygous ITS and confirmed by GISH and TBP. Shared introns between hybrids and the parental species are highlighted in Table S3. For the older tetraploid hybrid accessions NGY140 and NGY142 between *Le. aequinoctialis* and *Le. perpusilla*, the plastid sequences are different from both parents but closer to the latter. This suggests a long lasting, independent evolution of this lineage. This is also in agreement with some exclusive β-tubulin alleles, diverging from both parental species (*Le. aequinoctialis* and *Le. perpusilla*), but shared by all hybrid accessions in this cluster (see Supporting Information Table S3).

Among *Le. aequinoctialis s*.*s*. (blue dot), we distinguish three major subclusters, not always supported by high bootstrap values, but coincident with barcoding and ITS results. <u>Subcluster 1,</u> includes diploid *Le. aequinoctialis* accessions and a single triploid one (9648). According to TBP and ITS analyses, accession 9648 is heterozygous and merges alleles of diploid accessions from two branches of the same subcluster (Figs. **1** and **6**, Supporting Information Table S3), as result of an intraspecific cross. <u>Subcluster 2</u>, includes diploid and autotriploid accessions, because plastid, ITS and tubulin intron sequences are all from *Le. aequinoctialis*. <u>Subcluster</u> 3 exclusively comprises diploid accessions.

**Figure 6.**
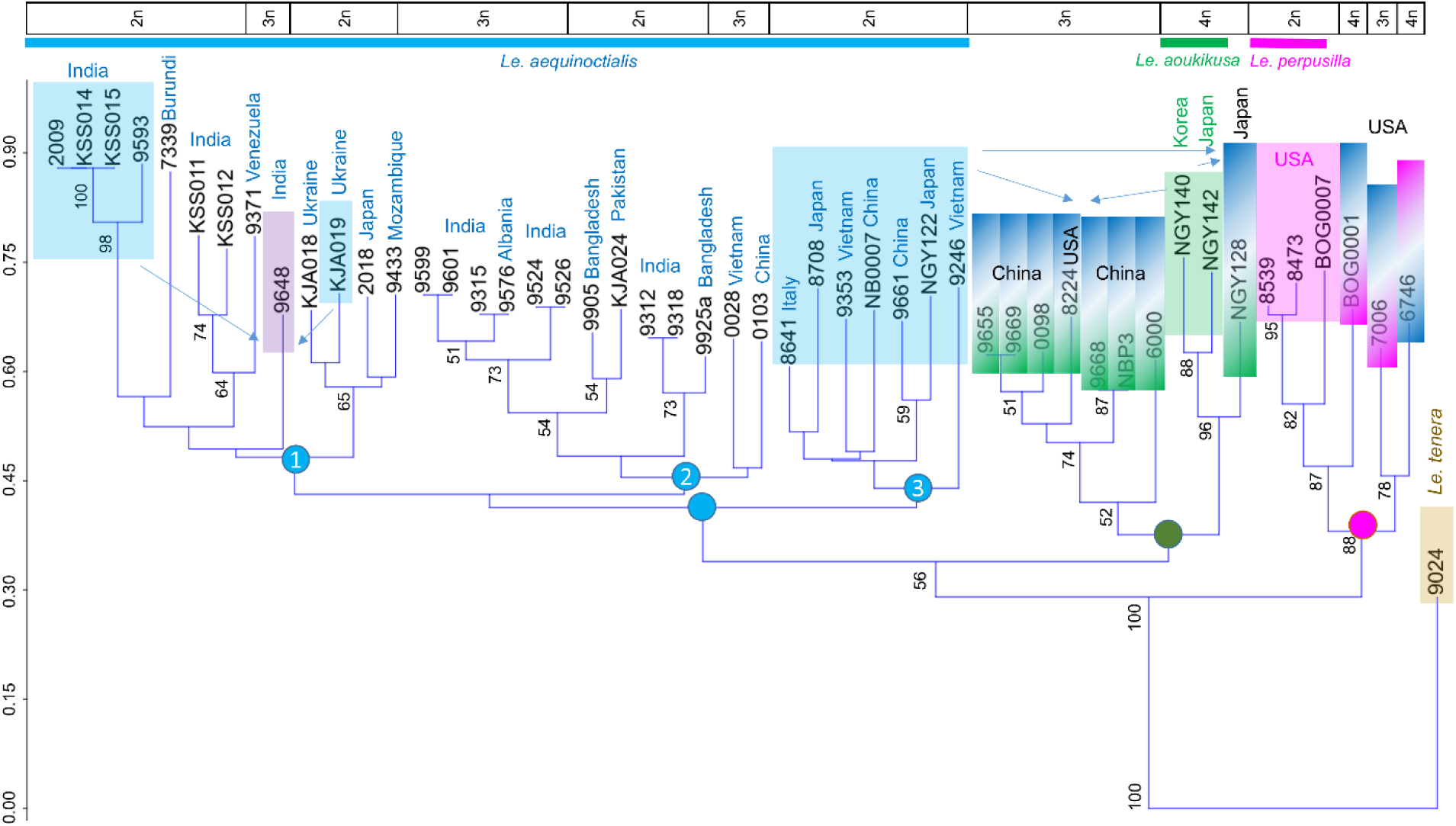
Phylogeny of presumed *Lemna aequinoctialis, Le. perpusilla, Le. aoukikusa* and *Le. tenera* (as an outgroup) accessions based on β-tubulin gene intron polymorphisms. Colored dots mark branches of *Le. perpusilla s.s.* and its recent hybrids with *Le. aequinoctialis* (pink dot), the ancient tetraploid hybrids between *Le. aequinoctialis* and *Le. perpusilla, Le.* × *aoukikusa,* and their triploid backcross hybrids with diploid *Le. aequinoctialis* (green dot), and *Le. aequinoctialis s.s.* and its autotriploid accessions and intraspecific hybrids (blue dot). In the upper part, ploidy level and geographic origin of the accessions are indicated. Arrows and/or color gradient indicate potential ancestors of hybrids; color gradients mark maternal (top) versus paternal (bottom) ancestors.

The allotriploid hybrids vary as to their genome size apparently depending on which *Le. aequinoctialis* and *Le. perpusilla* accession were the actual parents (Fig. **2**), For instance, 8224 (711 Mbp) could have obtained one genome of a paternal accession like 8473 (520 Mbp) and two maternal *Le. aequinoctialis* genomes of ∼450 Mbp. The differing genome sizes of the tetraploid hybrids are also explainable by combinations of correspondingly variable parental genomes. No polyploid hybrid has a larger genome size than expected from combination of potential parental accessions. This is reasonable, because interspecific hybridization results in loss rather than gain of genetic material.

## Discussion

### The evolution of the Alatae section

Our data demonstrate the reasons for the high variability as to genome size, chromosome number, morphological features and the discordance of phylogeny by chloroplast and nuclear markers among *Alatae* accessions. This variability is responsible for the difficulty to assign individual accessions to distinct taxa. Because of the relative easiness and frequency of fusion of reduced and/or unreduced gametes from the same or related, but different, taxa within the section *Alatae*, recurrent ploidy variants and hybrids appear in spite of the predominantly vegetative reproduction via sprouting of ‘daughter’ fronds from the meristematic pockets of the ‘mother’ fronds. Auto- and allotriploids, resulting from the merge of a reduced with an unreduced gamete, most likely propagate exclusively vegetatively. However, the allotetraploids retained (or re-gained) the ability to reproduce sexually and to hybridize further as suggested by the allotriploids of the *Le. aoukikusa* cluster (Fig. **6**), which apparently descended from a haploid female gamete of *Le. aequinoctialis* and a diploid male gamete of the allotetraploid hybrid *Le. aoukikusa*.

The allotetraploids 6746 and NGY142 were previously demonstrated to be self-fertile (Lee et al., 2024) but their hybrid nature was not known. So far, fertile allotetraploid duckweed hybrids are for the first time demonstrated here, but possibly also occur within the *Wolffia* genus. In contrast to the hitherto described hybrid species *Le. japonica* (Ernst et al., 2023, preprint) and *Le. mediterranea* (Braglia et al., 2024), some *Alatae* hybrids are tetraploid and fertile, while dihaploid hybrids were not yet found among *Alatae* accessions. As described for *Le. × mediterranea* (Braglia et al., 2024), multiple independent hybridizations have occurred also among *Alatae* species. The previously claimed species *Le. aoukikusa* (Lee et al., 2024) represents such a hybrid (see Figs. **1** and **6**). In addition, the iconic clone 6746, collected in 1954 in California and used for years in several experimental works, was first classified as *Le. perpusilla*,later emended to *Le. paucicostata* and finally ended up with the name *Le. aequinoctialis* (Landolt, 1986). Our demonstration of the hybrid origin of 6746, not only clarifies the reason of the uncertainty of its species assignment, but most likely explains also the reason for its homogamy and self-fertility which contrasts with the protogynous and self-sterile flowers of diploid *Le. aequinoctialis* (Lee et al., 2024). The same reproductive style is found in the *Le. aoukikusa* accession NGY142 (Lee et al., 2024), which is also a hybrid between *Le. aequinoctialis* and *Le. perpusilla*.

Because tri- and tetraploid *Alatae* hybrids occur in Northern America and in Asia as well, but there is not yet an unambiguous evidence for *Le. perpusilla* outside of Northern America, it seems reasonable to assume the interspecific hybrids occurred in Northern America and some of them successfully spread to Asia. Thus, the resolution of the complex *Alatae* section led to the following hypothetical scenario (see Fig. **7**).

**Figure 7.**
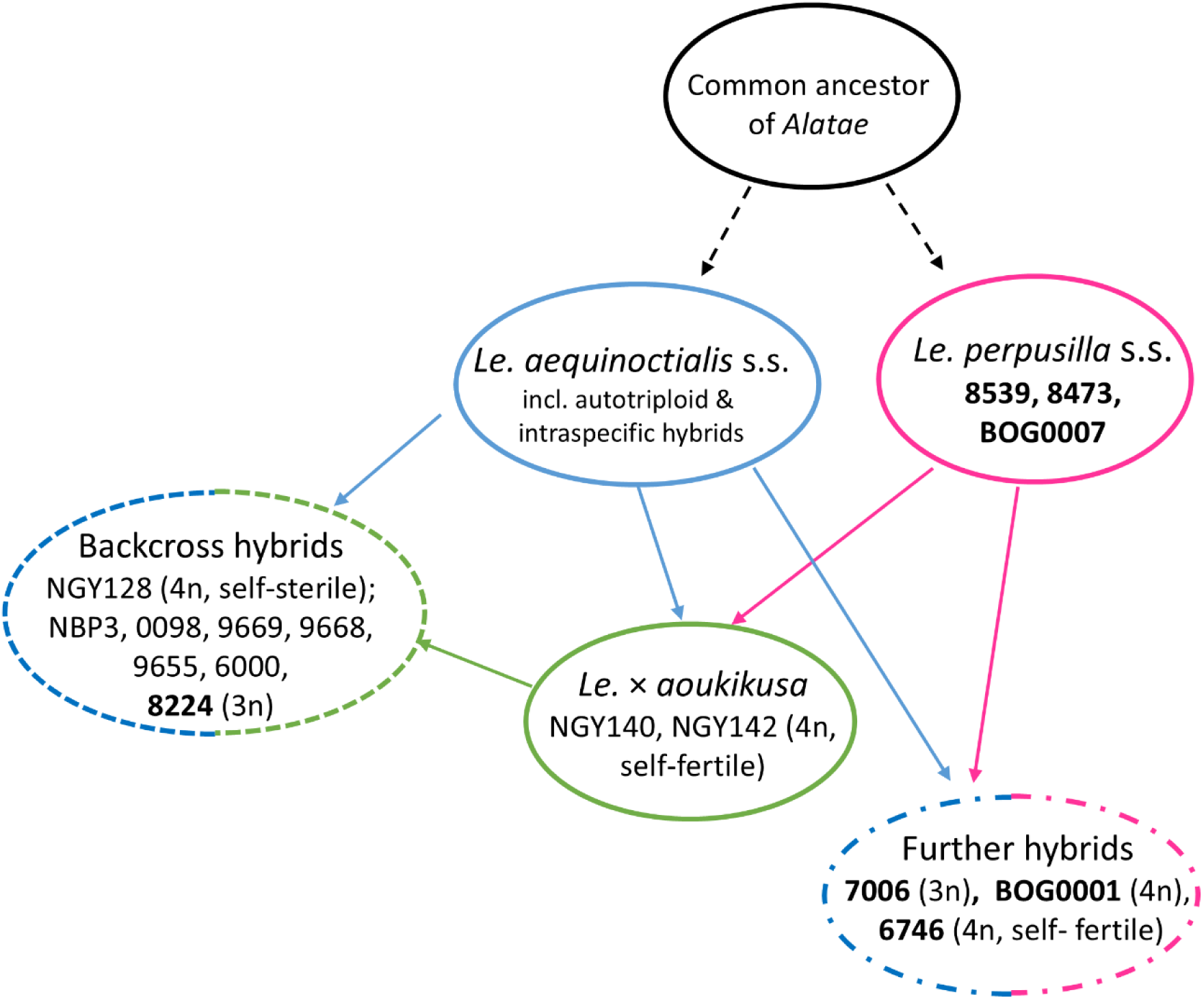
Hypothetical scenario of evolution within *Lemna* Section *Alatae.* In bold are accessions from America. The classically defined diploid species *Le. aequinoctialis* and *Le. perpusilla sensu stricto* were confirmed by the applied molecular methods. The previously named *Le. aoukikusa* accessions were uncovered as tetraploid interspecific hybrids between *Le. aequinoctialis* and *Le. perpusilla*. In addition, further tri- and tetraploid hybrids as well as tri- and tetraploid “Backcross hybrids” were detected from the USA and China, respectively.

Ancestors of the related and sexually proficient species *Le. aequinoctialis* and *Le. perpusilla,* with the ability to form unreduced gametes in addition to reduced ones, occurred together in the southern part of North America, where they hybridized repeatedly by fusion of unreduced gametes. The primary hybrid *Le. aoukikusa* migrated towards East-Asia due to its adaptation to lower temperatures. There, *Le. aoukikusa* itself backcrossed with *Le. aequinoctialis* (forming the allotriploid branch of the *Le. aoukikusa* cluster in Fig. **6**). One of the triploid hybrids, 8224, possibly migrated to USA. In parallel, autotriploids and intraspecific hybrids of *Le. aequinoctialis* arose in various geographic regions by fusion of reduced with unreduced gametes and spread there successfully. The reciprocal hybridization between *Le. aequinoctialis* and *Le. perpusilla*, forming tri- and tetraploid hybrids in North America (Fig. **6**) is apparently ongoing.

### Taxonomy of Alatae

An open question remains how to treat the *Alatae* accessions taxonomically. In addition to *Le. aequinoctialis s*.*s*. and *Le. perpusilla s*.*s*., we have the allotetraploid *Le. aoukikusa*, as well as a bunch of other tri- and tetraploid hybrids involving *Le. aequinoctialis* and *Le. perpusilla* or *Le. aoukikusa* (in the *Le. aoukikusa* and *Le. perpusilla* clusters, see Fig. **6**) for which the taxonomic status remains unclear. Although systematics of Lemnaceae has been extensively studied (Landolt, 1986; Les et al., 1997; Bog et al., 2019), only few considered the *Alatae* section, due to its complexity. Among them, recognition of *Le. perpusilla* as a separate species (Les et al., 1997) and the acceptance of *Le. aequinoctialis* as a correct name (Landolt, 1986; Sree et al., 2016). However, even now, this has not led to the termination of the use of the old nomenclature *Le. paucicostata* instead of the generally accepted one, which introduces additional confusion in taxonomic handling of *Alatae* accessions (16 references in https://pubmed.ncbi.nlm.nih.gov in the last 10 years). The variability of the morphological characteristics of *Alatae* accessions led to the use of different key criteria for species determination in different studies (Landolt, 1986; Sree et al., 2016; Bog et al., 2020a; Lee et al., 2024). Our molecular and cytogenetic analyses clearly confirmed two diploid species, but revealed distinct subgroups with a high level of variability among them. This correlates with previous studies indicating significant variation within the *Alatae* section (Tang et al., 2014; Tang et al., 2015; Chen et al., 2022; Lee et al., 2024). We also specified the genetic structure and geographic distribution of the previously presumed species *Le. aoukikusa* (Beppu & Takimoto, 1981; Lee et al., 2024) as an ancient tetraploid hybrid between *Le. aequinoctialis* and *Le. perpusilla* for which no diploid genotypes are known. Its self-fertility opens the possibility to consider it as a hybrid species, thus an evolving lineage, rather than a dead-end as are the dihaploid and triploid hybrids in the section *Lemna* (Romano et al., 2024 preprint). On the other hand, the highlighted complexity clearly indicates we are facing a taxonomically complex group (TCG; Ennos et al., 2005) in which the boundary of species blurs due to the presence of uniparental reproduction, high rate of inter-crossing, backcrossing, and polyploidization. This generates a genetically diverse mixture of related individuals, of more than one ploidy level, whose biological diversity defies simple classification into discrete species, according to the biological species concept (Ennos et al., 2005). Well studied examples are found in the *Senecio* and *Sorbus* genera (for reviews see Robertson et al., 2010; Wong et al., 2022). Assignment of a specimen to a species or a hybrid is possible only by a combination of approaches. Based on the available data we consider the *Alatae* accessions as *Le. aequinoctialis* species complex.

Summarizing, the data obtained allowed the first detailed assessment of the *Alatae* section from different continents using molecular and cytogenetic methods. A high level of diversity and the presence of ploidy variants and interspecific hybrids was demonstrated. The latter resulted from fusion of unreduced (or reduced and unreduced) gametes of different species and explains the taxonomic uncertainty within the section. Our results exemplify how the evolution of species of the genera *Wolffiella* and *Wolffia* of the duckweed family could be resolved in future.

## Author contribution

LM, IS, AS, LB, did conceptual work. AS, LB, JF, VS, PTNH, YL, SG, GC performed and evaluated experiments. AS, IS, LM, LB wrote the paper. All authors edited and approved the manuscript.

## Acknowledgements

We thank Klaus J. Appenroth, Nikolai Borisjuk, Manuela Bog, Sowjanya Sree and Eric Lam for providing accessions and KJA for critical reading of the manuscript. AS thanks the German National Academy of Sciences Leopoldina for a Leopoldina Ukraine Distinguished Fellowship which made these investigations possible. Part of this study was carried out within the Agritech National Research Center and received funding from the European Union Next-Generation EU [Piano Nazionale Di Ripresa e Resilienza (PNRR), Missione 4 Componente 2, Investimento 1.4— Project CN00000022] to the National Research Council. This manuscript reflects only the authors’ views and opinions; neither the European Union nor the European Commission can be considered responsible for them.

## Supporting Information (brief legends)

**Table S1** The list of duckweed accessions and research conducted on them

**Table S2** List of primers used in this study.

**Table S3** Schematic representation of the numerical output of TBP profiles for each analyzed accession

**Fig. S1** Alignment of the *atpF*-*atpH* spacers of *Le. aequinoctialis*, *Le. aoukikusa*, *Le. perpusilla* and *Le. tenera* clones.

**Fig. S2** Alignment of *psbK*–*psbI* spacers of *Le. aequinoctialis*, *Le. aoukikusa*, *Le. Perpusilla* and *Le. tenera* clones.

**Fig. S3** Alignment of ITS1-5.8S-ITS2 sequences of *Le. aequinoctialis*, *Le. aoukikusa*, *Le. perpusilla* and *Le. tenera* clones.

