## Supporting Information Figure S1-S3 for "Genome diversity and phylogeny of the section *Alatae* of genus *Lemna* (Lemnaceae), comprising the presumed species *Lemna aequinoctialis, Le. perpusilla* and *Le. aoukikusa*"

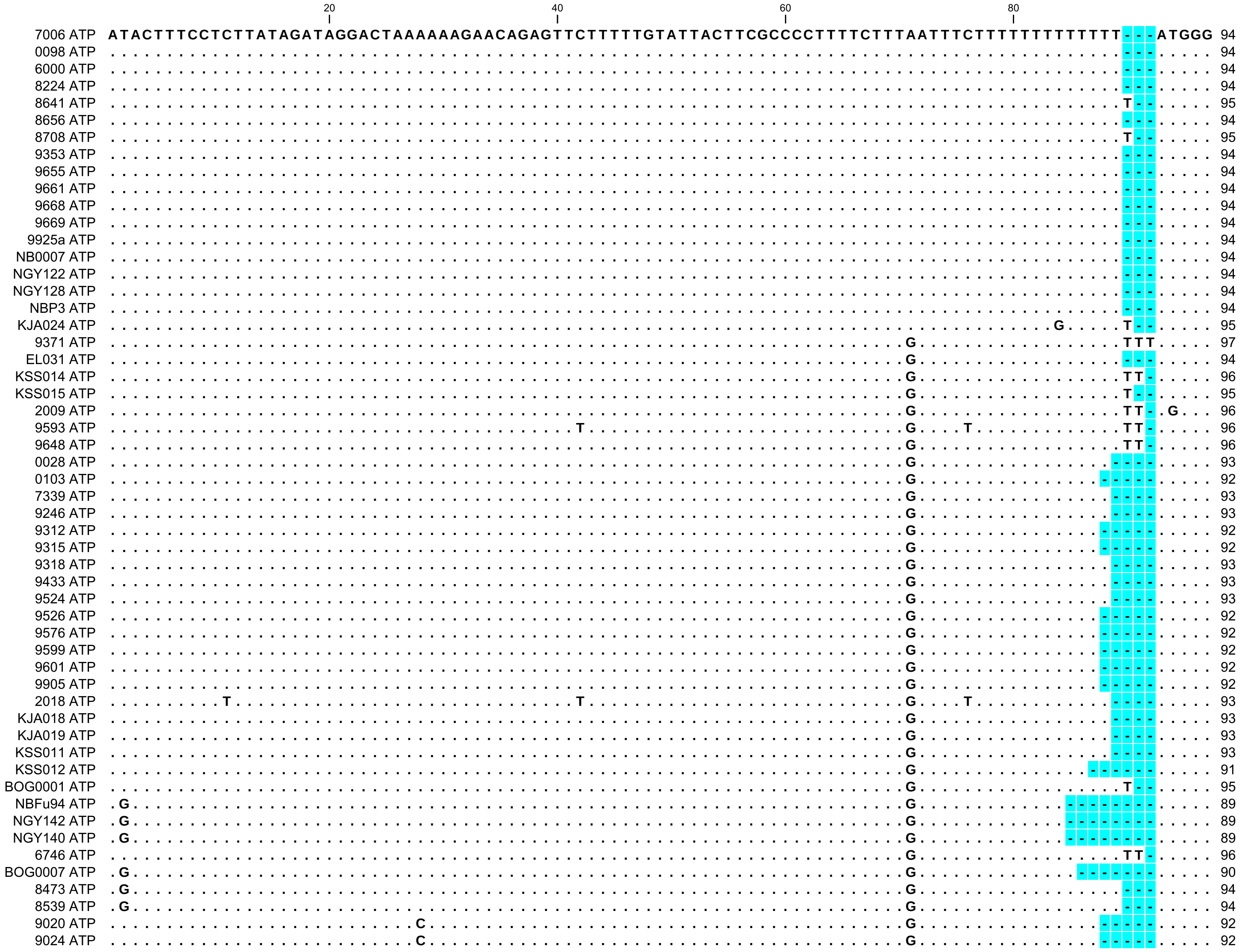

Figure S1 Alignment of the *atpF*-*atpH* spacers of *Le. aequinoctialis*, *Le. aoukikusa*, *Le. perpusilla* and *Le. tenera* clones. Matching residues are shown as dots. Gaps are highlighted in light-blue.

```
               100                 120                 140                 160                 180
    7006 ATP ATTTTTAAATAGAATAGATGACATTAATTAACTTAATT-----GAGAACTTTTTTATTTATTATTTTATTCTAATTAAAGTTTACAATTACAAGAGC 186
    0098 ATP ......................................-----...................................................... 186
    6000 ATP ......................................-----...................................................... 186
    8224 ATP ......................................-----...................................................... 186
    8641 ATP ......................................-----...................................................... 187
    8656 ATP ......................................-----...................................................... 186
    8708 ATP ......................................-----...................................................... 187
    9353 ATP ......................................-----...................................................... 186
    9655 ATP ......................................-----...................................................... 186
    9661 ATP ......................................-----...................................................... 186
    9668 ATP ......................................-----...................................................... 186
    9669 ATP ......................................-----...................................................... 186
   9925a ATP ......................................-----...................................................... 186
  NB0007 ATP ......................................TAATT...................................................... 191
  NGY122 ATP ......................................TAATT...................................................... 191
  NGY128 ATP ......................................TAATT...................................................... 191
    NBP3 ATP ......................................-----...................................................... 186
  KJA024 ATP .....A................................-----...................................................... 187
    9371 ATP .....A................................-----...................................................... 189
   EL031 ATP .....A................................-----...................................................... 186
  KSS014 ATP .....A................................TAATT...................................................... 193
  KSS015 ATP .....A................................TAATT...................................................... 192
    2009 ATP .....A................................TAATT...................................................... 193
    9593 ATP .....A................................TAATT...................................................... 193
    9648 ATP .....A.....A..........................TAATT..A................................................... 193
    0028 ATP .....A................................-----...................................................... 185
    0103 ATP .....A................................-----...................................................... 184
    7339 ATP .....A................................-----...................................................... 185
    9246 ATP .....A................................-----...................................................... 185
    9312 ATP .....A................................-----...................................................... 184
    9315 ATP .....A................................-----...................................................... 184
    9318 ATP .....A................................-----...................................................... 185
    9433 ATP .....A................................-----...................................................... 185
    9524 ATP .....A................................-----...................................................... 185
    9526 ATP .....A................................-----...................................................... 184
    9576 ATP .....A................................-----...................................................... 184
    9599 ATP .....A................................-----...................................................... 184
    9601 ATP .....A................................-----...................................................... 184
    9905 ATP .....A................................-----...................................................... 184
    2018 ATP .....A................................-----...................................................... 185
  KJA018 ATP .....A................................-----...................................................... 185
  KJA019 ATP .....A................................-----...................................................... 185
  KSS011 ATP .....A................................-----...................................................... 185
  KSS012 ATP .....A................................-----...................................................... 183
 BOG0001 ATP .....A................................-----...................................................... 187
  NBFu94 ATP .....A....G...........................-----...................................................... 181
  NGY142 ATP .....A....G...........................-----...................................................... 181
  NGY140 ATP .....A....G...........................-----...................................................... 181
    6746 ATP .....A....G...........................-----...................................................... 188
 BOG0007 ATP .....A....G...........................-----...................................................... 182
    8473 ATP .....A....G...........................-----...................................................... 186
    8539 ATP .....A....G...........................-----...................................................... 186
    9020 ATP .....A....G...........................-----...................................................... 184
    9024 ATP .....A....G...........................-----...................................................... 184
```

```
                    200                 220                 240                 260                 280
                    |                   |                   |                   |                   |
7006 ATP     ATACTTATTGGGTTAGGGCCTGACTATTTTGTCAATAAATACCTTGTTTGTTGCGTTACAACGCATACTCAAAAAAGTTTTCCCTTACATTATACTA 283
0098 ATP     ................................................................................................. 283
6000 ATP     ................................................................................................. 283
8224 ATP     ................................................................................................. 283
8641 ATP     ................................................................................................. 284
8656 ATP     ................................................................................................. 283
8708 ATP     ................................................................................................. 284
9353 ATP     ................................................................................................. 283
9655 ATP     ................................................................................................. 283
9661 ATP     ................................................................................................. 283
9668 ATP     ................................................................................................. 283
9669 ATP     ................................................................................................. 283
9925a ATP    ................................................................................................. 283
NB0007 ATP   ................................................................................................. 288
NGY122 ATP   ................................................................................................. 288
NGY128 ATP   ................................................................................................. 288
EL031 ATP    ................................................................................................. 283
KSS014 ATP   ................................................................................................. 290
KSS015 ATP   ................................................................................................. 289
2009 ATP     ................................................................................................. 290
9593 ATP     ................................................................................................. 290
9648 ATP     ................................................................................................. 290
0028 ATP     ..........................................................G...................................... 282
0103 ATP     ..........................................................G...................................... 281
7339 ATP     ..........................................................G...................................... 282
9246 ATP     ..........................................................G...................................... 282
9312 ATP     ..........................................................G...................................... 281
9315 ATP     ..........................................................G...................................... 281
9318 ATP     ..........................................................G...................................... 282
9433 ATP     ..........................................................G...................................... 282
9524 ATP     ..........................................................G...................................... 282
9526 ATP     ..........................................................G...................................... 281
9576 ATP     ..........................................................G...................................... 281
9599 ATP     ..........................................................G...................................... 281
9601 ATP     ..........................................................G...................................... 281
9905 ATP     ..........................................................G...................................... 281
2018 ATP     ..........................................................G...................................... 282
KJA018 ATP   ..........................................................G...................................... 282
KJA019 ATP   ..........................................................G...................................... 282
KSS011 ATP   ..........................................................G...................................... 282
KSS012 ATP   ..........................................................G...................................... 280
BOG0001 ATP  ................................................................................................. 284
NBFu94 ATP   ...................................................................................A............. 278
NGY142 ATP   ...................................................................................A............. 278
NGY140 ATP   ...................................................................................A............. 278
6746 ATP     ...................................................................................A............. 285
BOG0007 ATP  ...................................................................................A............. 279
8473 ATP     ..............................A....................................................A............. 283
8539 ATP     ..............................A....................................................A............. 283
9020 ATP     .................T.......G...................A.....................................A............. 281
9024 ATP     .................T.......G...................A.....................................A............. 281
```

```
              300                 320                 340                 360                 380
7006 ATP      AGAACTAAAAACGGGAAGGAAGAAAGCGAGAGGATCTGCTAATTACTAATCCTAAAATCAGTCCTTCCCGGAGGTATTCTCTCAACGAATAAGTAAT 380
0098 ATP      ................................................................................................. 380
6000 ATP      ................................................................................................. 380
8224 ATP      ................................................................................................. 380
8641 ATP      ................................................................................................. 381
8656 ATP      ................................................................................................. 380
8708 ATP      ................................................................................................. 381
9353 ATP      ................................................................................................. 380
9655 ATP      ................................................................................................. 380
9661 ATP      ................................................................................................. 380
9668 ATP      ................................................................................................. 380
9669 ATP      ................................................................................................. 380
9925a ATP     ................................................................................................. 380
NB0007 ATP    ................................................................................................. 385
NGY122 ATP    ................................................................................................. 385
NGY128 ATP    ................................................................................................. 385
EL031 ATP     ................................................................................................. 380
KSS014 ATP    ................................................................................................. 387
KSS015 ATP    ................................................................................................. 386
2009 ATP      ................................................................................................. 387
9593 ATP      ................................................................................................. 387
9648 ATP      ................................................................................................. 387
0028 ATP      ................................................................................................. 379
0103 ATP      ................................................................................................. 378
7339 ATP      ................................................................................................. 379
9246 ATP      ................................................................................................. 379
9312 ATP      ................................................................................................. 378
9315 ATP      ................................................................................................. 378
9318 ATP      ................................................................................................. 379
9433 ATP      ................................................................................................. 379
9524 ATP      ................................................................................................. 379
9526 ATP      ................................................................................................. 378
9576 ATP      ................................................................................................. 378
9599 ATP      ................................................................................................. 378
9601 ATP      ................................................................................................. 378
9905 ATP      ................................................................................................. 378
2018 ATP      ................................................................................................. 379
KJA018 ATP    ................................................................................................. 379
KJA019 ATP    ......................................................................................A.......... 379
KSS011 ATP    ................................................................................................. 379
KSS012 ATP    ................................................................................................. 377
BOG0001 ATP   ................................................................................................. 381
BOG0007 ATP   .....A........................................................................................... 376
8473 ATP      ................................................................................................. 380
8539 ATP      ................................................................................................. 380
9020 ATP      ................................................................................................. 378
9024 ATP      ................................................................................................. 378
```

```
                       400                 420                 440                 460
                       |                   |                   |                   |
7006 ATP      TGTTAGAGTGCAATGTTGATAGAATTCGAAGAAGCAAAAAGCAAGTCTAAGTCAAAAAGT---------ACTTTCTTTTTTGTAGAA 458
0098 ATP      ............................................................---------.................. 458
6000 ATP      ............................................................---------.................. 458
8224 ATP      ............................................................---------.................. 458
8641 ATP      ............................................................---------.................. 459
8656 ATP      ............................................................---------.................. 458
8708 ATP      ............................................................---------.................. 459
9353 ATP      ............................................................---------.................. 458
9655 ATP      ............................................................---------.................. 458
9661 ATP      ............................................................---------.................. 458
9668 ATP      ............................................................---------.................. 458
9669 ATP      ............................................................---------.................. 458
9925a ATP     ............................................................---------.................. 458
NB0007 ATP    ............................................................---------.................. 463
NGY122 ATP    ............................................................---------.................. 463
NGY128 ATP    ............................................................---------.................. 463
NBP3 ATP      ............................................................---------.................. 458
KJA024 ATP    ............................................................---------.................. 459
9371 ATP      ............................................................---------.................. 461
EL031 ATP     ............................................................---------.................. 458
KSS014 ATP    ............................................................---------.................. 465
KSS015 ATP    ............................................................---------.................. 464
2009 ATP      ............................................................---------.................. 465
9593 ATP      ............................................................---------.................. 465
9648 ATP      ............................................................---------.................. 465
0028 ATP      ............................................................---------.................. 457
0103 ATP      ............................................................---------.................. 456
7339 ATP      ............................................................---------.................. 457
9246 ATP      ............................................................---------.................. 457
9312 ATP      ............................................................---------.................. 456
9315 ATP      ............................................................---------.................. 456
9318 ATP      ............................................................---------.................. 457
9433 ATP      ............................................................---------.................. 457
9524 ATP      ............................................................---------.................. 457
9526 ATP      ............................................................---------.................. 456
9576 ATP      ............................................................---------.................. 456
9599 ATP      ............................................................---------.................. 456
9601 ATP      ............................................................---------.................. 456
9905 ATP      ............................................................---------.................. 456
2018 ATP      ............................................................---------.................. 457
KJA018 ATP    ............................................................---------.................. 457
KJA019 ATP    ............................................................---------.................. 457
KSS011 ATP    ............................................................---------.................. 457
KSS012 ATP    ............................................................---------.................. 455
BOG0001 ATP   ............................................................---------.................. 459
NBFu94 ATP    ............................................................---------...........-...... 452
NGY142 ATP    ............................................................---------...........-...... 452
NGY140 ATP    ............................................................---------...........-...... 452
6746 ATP      ............................................................---------...........-...... 459
BOG0007 ATP   ............................................................---------...........-...... 453
8473 ATP      ............................................................---------...........-...... 457
8539 ATP      ............................................................---------...........-...... 457
9020 ATP      .................................T..........................CTATTACGT.................. 465
9024 ATP      .................................T..........................CTATTACGT.................. 465
```

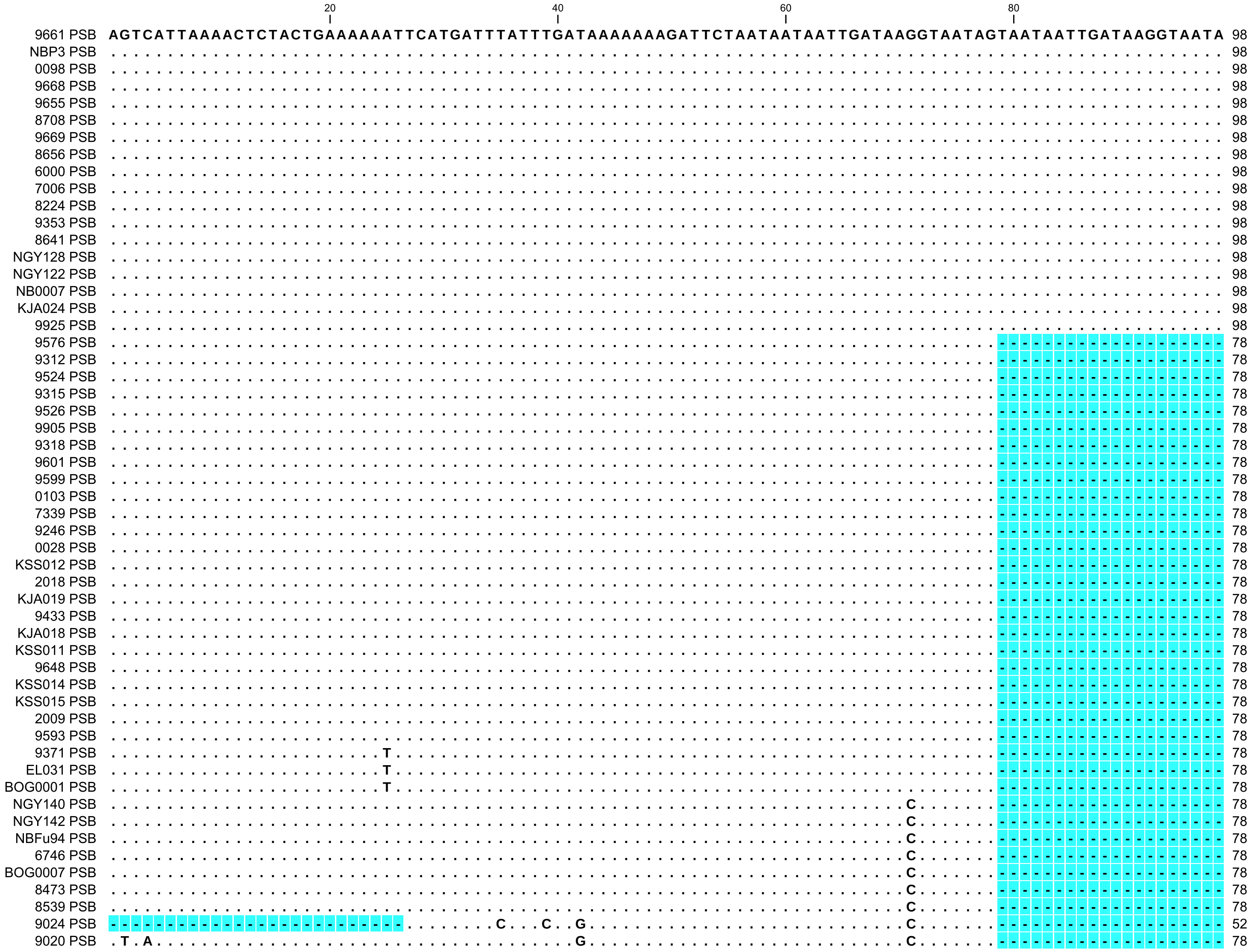

Figure S2 Alignment of *psbK–psbI* spacers of *Le. aequinoctialis*, *Le. aoukikusa*, *Le. perpusilla* and *Le. tenera* clones. Matching residues are shown as dots. Gaps are highlighted in light-blue.

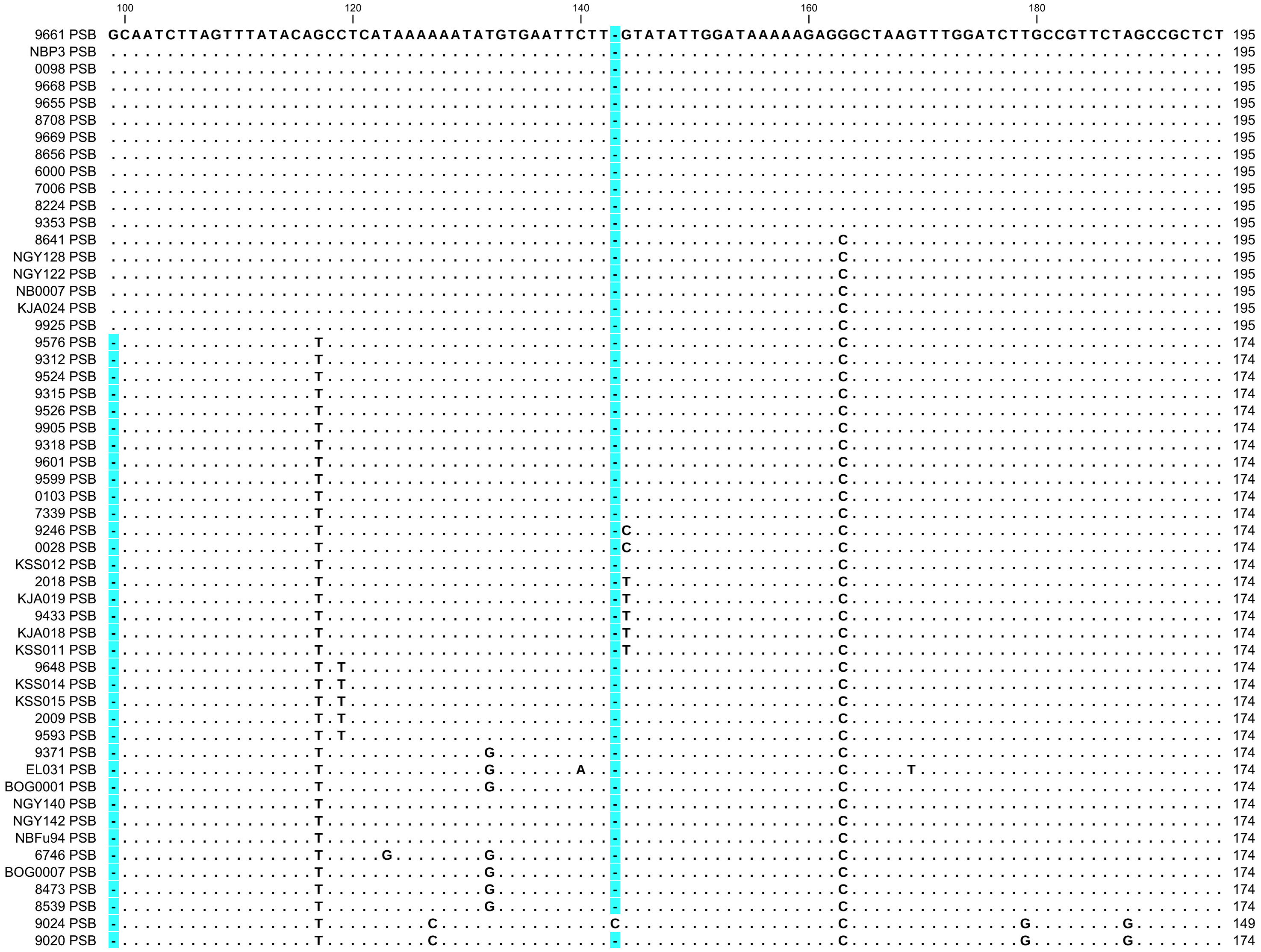

```
               200                 220                 240                 260                 280
9661 PSB    TCCAGTGAGTAACTACTTTATTAGCTTTTGTTTTACACAAAACTTTATTGTTATATTATATTATTAGAGTTAATAGTCGAATAACCTTTTGGATAATG 293
NBP3 PSB    .................................................................................................. 293
0098 PSB    .................................................................................................. 293
9668 PSB    .................................................................................................. 293
9655 PSB    .................................................................................................. 293
8708 PSB    .................................................................................................. 293
9669 PSB    .................................................................................................. 293
8656 PSB    .................................................................................................. 293
6000 PSB    .................................................................................................. 293
7006 PSB    .................................................................................................. 293
8224 PSB    .................................................................................................. 293
9353 PSB    .................................................................................................. 293
8641 PSB    .................................................................................................. 293
NGY128 PSB .................................................................................................. 293
NGY122 PSB .................................................................................................. 293
NB0007 PSB .................................................................................................. 293
KJA024 PSB .............................T.................................................................... 293
9925 PSB    .............................T.................................................................... 293
9576 PSB    ...........................................................-----.................................. 267
9312 PSB    ...........................................................-----.................................. 267
9524 PSB    ...........................................................-----.................................. 267
9315 PSB    ...........................................................-----.................................. 267
9526 PSB    ...........................................................-----.................................. 267
9905 PSB    ...........................................................-----.................................. 267
9318 PSB    ...........................................................-----.................................. 267
9601 PSB    ...........................................................-----.................................. 267
9599 PSB    ...........................................................-----.................................. 267
0103 PSB    ...........................................................-----.................................. 267
7339 PSB    ...........................................................-----.................................. 267
9246 PSB    ...........................................................-----.................................. 267
0028 PSB    ...........................................................-----.................................. 267
KSS012 PSB ...........................................................-----.................................. 267
2018 PSB    ...........................................................-----.................................. 267
KJA019 PSB ...........................................................-----.................................. 267
9433 PSB    ...........................................................-----.................................. 267
KJA018 PSB ...........................................................-----.................................. 267
KSS011 PSB ...........................................................-----.................................. 267
9648 PSB    .A.........................................................-----.................................. 267
KSS014 PSB .A.........................................................-----.................................. 267
KSS015 PSB .A.........................................................-----.................................. 267
2009 PSB    .A.........................................................-----.................................. 267
9593 PSB    .A.........................................................-----.................................. 267
9371 PSB    ...........................................................-----.................................. 267
EL031 PSB   ...........................................................-----.................................. 267
BOG0001 PSB ...........................................................-----.................................. 267
NGY140 PSB ...........................................................-----.................................. 267
NGY142 PSB ...........................................................-----.................................. 267
NBFu94 PSB ...........................................................-----.................................. 267
6746 PSB    ...........................................................-----.................................. 267
BOG0007 PSB .........................................................T.-----.................................C 267
8473 PSB    ...........................................................-----.................................. 267
8539 PSB    ...........................................................-----.................................. 267
9024 PSB    ...................C..................................-------..............T.........A....C......C 240
9020 PSB    ...................C..................................-------..............T.........A....C......C 265
```

```
             300                 320                 340                 360                 380
9661 PSB     AACAAGTTATAATCTTAATTCAAAAAATTCATGAATTTGAAAATTCAGTTTTTCTA-----GAAAAAAACACTGAATACTTAATTCAATTAAAAAACA 386
NBP3 PSB     ........................................................-----..................................... 386
0098 PSB     ........................................................-----..................................... 386
9668 PSB     ........................................................-----..................................... 386
9655 PSB     ........................................................-----..................................... 386
8708 PSB     ........................................................-----..................................... 386
9669 PSB     ........................................................-----..................................... 386
8656 PSB     ........................................................-----..................................... 386
6000 PSB     ........................................................-----..................................... 386
7006 PSB     ........................................................-----..................................... 386
8224 PSB     ........................................................-----..................................... 386
9353 PSB     ........................................................-----..................................... 386
8641 PSB     ........................................................-----..................................... 386
NGY128 PSB   ........................................................-----..................................... 386
NGY122 PSB   ........................................................-----..................................... 386
NB0007 PSB   ........................................................-----..................................... 386
KJA024 PSB   ........................................................-----..................................... 386
9925 PSB     ........................................................-----..................................... 386
9576 PSB     ........................................................-----..................................... 360
9312 PSB     ........................................................-----..................................... 360
9524 PSB     ........................................................-----..................................... 360
9315 PSB     ........................................................-----..................................... 360
9526 PSB     ........................................................-----..................................... 360
9905 PSB     ........................................................-----..................................... 360
9318 PSB     ........................................................-----..................................... 360
9601 PSB     ........................................................-----..................................... 360
9599 PSB     ........................................................-----..................................... 360
0103 PSB     ........................................................-----..................................... 360
7339 PSB     ........................................................-----..................................... 360
9246 PSB     ........................................................-----..................................... 360
0028 PSB     ........................................................-----..................................... 360
KSS012 PSB   ........................................................-----..................................... 360
2018 PSB     ......................................T.................-----.......-............................. 359
KJA019 PSB   ......................................T.................-----.......-............................. 359
9433 PSB     ......................................T.................-----.......-............................. 359
KJA018 PSB   ......................................T.................-----.......-............................. 359
KSS011 PSB   ......................................T.................-----.......-............................. 359
9648 PSB     ........................................................-----..................................... 360
KSS014 PSB   ........................................................-----..................................... 360
KSS015 PSB   ........................................................-----..................................... 360
2009 PSB     ........................................................-----..................................... 360
9593 PSB     ........................................................-----..................................... 360
9371 PSB     ......................C.................................-----..................................... 360
EL031 PSB    ......................C.................................-----..................................... 360
BOG0001 PSB  ......................C.................................-----..................................... 360
NGY140 PSB   ..G.....................................................-----..................................... 360
NGY142 PSB   ..G.....................................................-----..................................... 360
NBFu94 PSB   ..G.....................................................-----..................................... 360
6746 PSB     ........................................................-----......--............................. 358
BOG0007 PSB  ........................................................-----.......-............................. 359
8473 PSB     ........................................................-----.......-............................. 359
8539 PSB     ........................................................-----.......-............................. 359
9024 PSB     .C......................................................TACTA..............C..T.................A. 338
9020 PSB     .C......................................................TACTA..............C..T.................A. 363
```

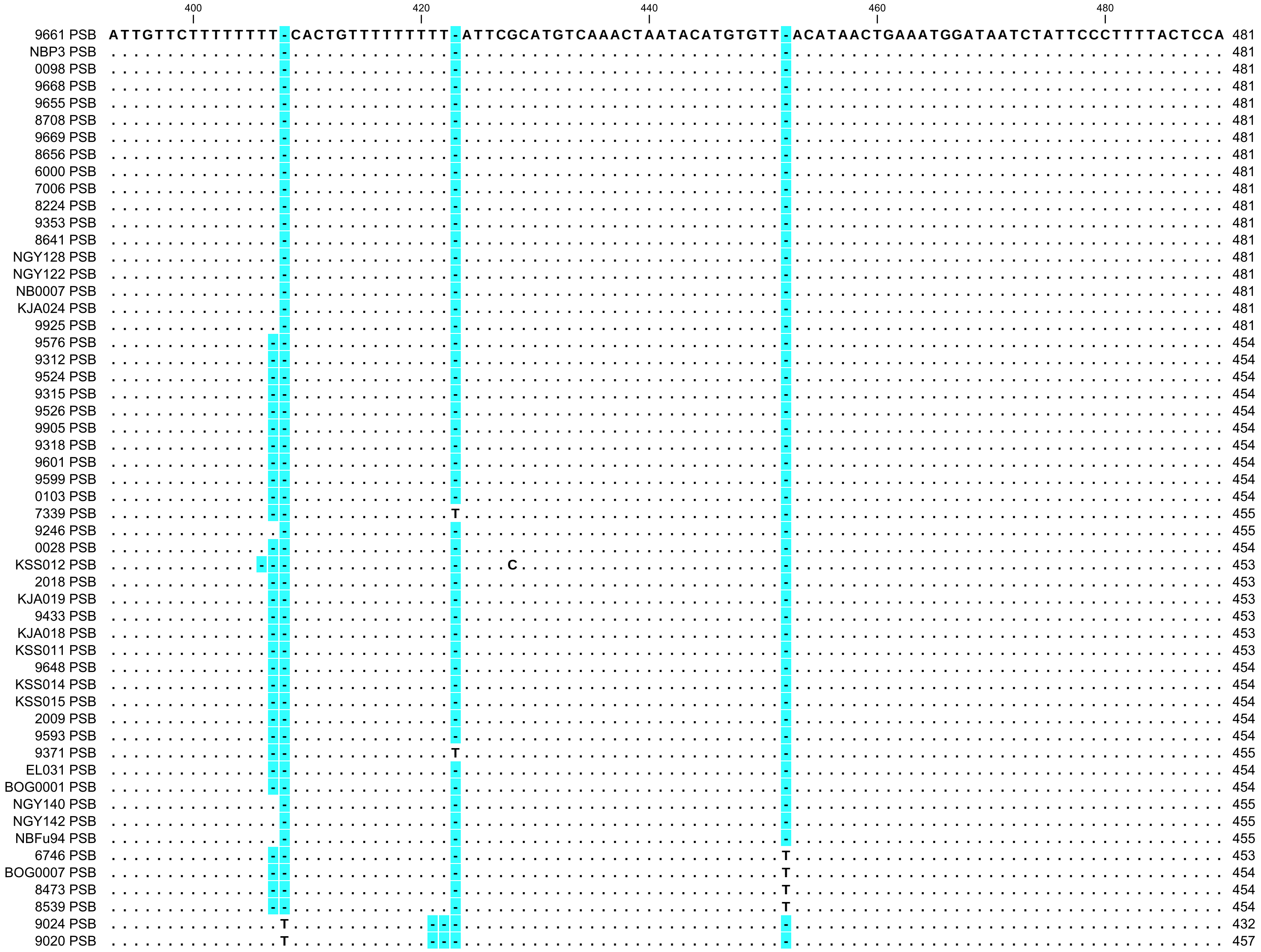
400
420
440
460
480
9661 PSB ATTGTTCTTTTTTTT-CACTGTTTTTTTTT-ATTCGCATGTCAAACTAATACATGTGTT-ACATAACTGAAATGGATAATCTATTCCCTTTTACTCCA 481
NBP3 PSB 481
0098 PSB 481
9668 PSB 481
9655 PSB 481
8708 PSB 481
9669 PSB 481
8656 PSB 481
6000 PSB 481
7006 PSB 481
8224 PSB 481
9353 PSB 481
8641 PSB 481
NGY128 PSB 481
NGY122 PSB 481
NB0007 PSB 481
KJA024 PSB 481
9925 PSB 481
9576 PSB 454
9312 PSB 454
9524 PSB 454
9315 PSB 454
9526 PSB 454
9905 PSB 454
9318 PSB 454
9601 PSB 454
9599 PSB 454
0103 PSB 454
7339 PSB 455
9246 PSB 455
0028 PSB 454
KSS012 PSB 453
2018 PSB 453
KJA019 PSB 453
9433 PSB 453
KJA018 PSB 453
KSS011 PSB 453
9648 PSB 454
KSS014 PSB 454
KSS015 PSB 454
2009 PSB 454
9593 PSB 454
9371 PSB 455
EL031 PSB 454
BOG0001 PSB 454
NGY140 PSB 455
NGY142 PSB 455
NBFu94 PSB 455
6746 PSB 453
BOG0007 PSB 454
8473 PSB 454
8539 PSB 454
9024 PSB 432
9020 PSB 457

```
                     500
                      |
9661 PSB     AAAA-TGATCCAATCTTGGAGATTGTGTA 509
NBP3 PSB     ....-........................ 509
0098 PSB     ....-........................ 509
9668 PSB     ....-........................ 509
9655 PSB     ....-........................ 509
8708 PSB     ....-........................ 509
9669 PSB     ....-........................ 509
8656 PSB     ....-........................ 509
6000 PSB     ....-........................ 509
7006 PSB     ....-........................ 509
8224 PSB     ....-........................ 509
9353 PSB     ....-........................ 509
8641 PSB     ....-........................ 509
NGY128 PSB   ....-........................ 509
NGY122 PSB   ....-........................ 509
NB0007 PSB   ....-........................ 509
KJA024 PSB   ....-........................ 509
9925 PSB     ....-........................ 509
9576 PSB     ....-........................ 482
9312 PSB     ....-........................ 482
9524 PSB     ....-........................ 482
9315 PSB     ....-........................ 482
9526 PSB     ....-........................ 482
9905 PSB     ....-........................ 482
9318 PSB     ....-........................ 482
9601 PSB     ....-........................ 482
9599 PSB     ....-........................ 482
0103 PSB     ....-........................ 482
7339 PSB     ....-........................ 483
9246 PSB     ....-........................ 483
0028 PSB     ....-........................ 482
KSS012 PSB   ....-........................ 481
2018 PSB     ....-........................ 481
KJA019 PSB   ....-........................ 481
9433 PSB     ....-........................ 481
KJA018 PSB   ....-........................ 481
KSS011 PSB   ....-........................ 481
9648 PSB     ....-........................ 482
KSS014 PSB   ....-........................ 482
KSS015 PSB   ....-........................ 482
2009 PSB     ....-........................ 482
9593 PSB     ....-........................ 482
9371 PSB     ....-........................ 483
EL031 PSB    ....-........................ 482
BOG0001 PSB  ....-........................ 482
NGY140 PSB   ....-........................ 483
NGY142 PSB   ....-........................ 483
NBFu94 PSB   ....-........................ 483
6746 PSB     ....-........................ 481
BOG0007 PSB  ....-........................ 482
8473 PSB     ....-........................ 482
8539 PSB     ....-........................ 482
9024 PSB     ....A........................ 461
9020 PSB     ....A........................ 486
```

```
                                    20                  40                  60                  80                  100
0098      TCGTCGAAACCCCGGCCCCGGCCGGCGGACGCACGAGAGAACGGTCTACCGAGCGTGGCACCCGCCCGGTTCCGCGCGCCC-GTCGCGGCGGCCGCTCCG 99
NGY128-1  .................................................................................-.................. 99
7006      .................................................................................-.................. 99
8708      .................................................................................-.................. 99
9246      .................................................................................-.................. 99
9312      .................................................................................-.................. 99
9315      .................................................................................-.................. 99
9318      .........A.......................................................................-.................. 99
9524      .................................................................................-.................. 99
9576      .................................................................................-.................. 99
9599      .................................................................................-.................. 99
9601      .................................................................................-.................. 99
9648-1    .................................................................................-.................. 99
9655      .................................................................................-.................. 99
9661      .................................................................................-.................. 99
9668      .................................................................................-.................. 99
9669      .................................................................................-.................. 99
9905      .................................................................................-.................. 99
9925a     .................................................................................-.................. 99
NBFu94    .................................................................................-.................. 99
KSS011    .................................................................................-.................. 99
NB0007    .................................................................................-.................. 99
NBP3      .................................................................................-.................. 99
8224      .................................................................................-.................. 99
0028      .................................................................................-.................. 99
0103      .................................................................................-.................. 99
6000      .................................................................................-.................. 99
8641      .................................................................................-.................. 99
NGY122    .................................................................................-.................. 99
2018      ......................................................................C..........-.................. 99
9433      ......................................................................C..........-.................. 99
KJA019    ......................................................................C..........-.................. 99
8656      .................................................................................-.................. 99
9526      .................................................................................-.................. 99
KJA024    .................................................................................-.................. 99
KJA018    ......................................................................C..........-.................. 99
9353      .................................................................................-.................. 99
2009      ..............................A.........................C........................-.................. 99
9648-2    ..............................A.........................C........................-.................. 99
KSS014    ..............................A.........................C........................-.................. 99
KSS015    ..............................A.........................C........................-.................. 99
9593      ........................................................C........................-.................. 99
KSS012    ..............................A.........................C........................-.................. 99
7339      .....................................C..................C........................-.................. 99
6746      .....................................C....A......................................-.................C 99
9371      .....................................C........................................G..-.................. 99
EL031     .....................................C..................C........................-..........G....... 99
NGY140    ......C..............................C..................C....G.A.................C.A................ 100
NGY142    ......C..............................C..................C....G.A.................C.A................ 100
NGY128-2  ......C..............................C..................C....G.A.................C.A................ 100
8473      ......C..............................C..................C....ATA.................C.A................ 100
8539      ......C..............................C..................C....ATA.................C.A................ 100
BOG0007   ......C..............................C..................C....ATA.................C.A................ 100
BOG0001   ......C...T..........................C..................C....ATA.................C.A................ 100
9020      ......C..............................C..................C....GGAG..............G.C.A........-------- 92
9024      ......C..............................C..................C....GGAG..............G.C.A........-------- 92
```

Figure S3 Alignment of ITS1-5.8S-ITS2 sequences of *Le. aequinoctialis*, *Le. aoukikusa*, *Le. perpusilla* and *Le. tenera* clones. Matching residues are shown as dots. Gaps are highlighted in light-blue.

```
                120                 140                 160                 180                 200
0098       AGCGCCCGGCGGGCTCCCCGCGCGCGCCCTCGGGCGCGCCGGCGGGGCCCCCGGGCCGCGGGGCGTCCCCCGCGGCGGTCGGGCGGGCCGGACA--ACCG 197
NGY128-1   ............................................................................................--.... 197
7006       ............................................................................................--.... 197
8708       ............................................................................................--.... 197
9246       ............................................................................................--.... 197
9312       ............................................................................................--.... 197
9315       ............................................................................................--.... 197
9318       ............................................................................................--.... 197
9524       ............................................................................................--.... 197
9576       ............................................................................................--.... 197
9599       ............................................................................................--.... 197
9601       ............................................................................................--.... 197
9648-1     ............................................................................................--.... 197
9655       ............................................................................................--.... 197
9661       ............................................................................................--.... 197
9668       ............................................................................................--.... 197
9669       ............................................................................................--.... 197
9905       ............................................................................................--.... 197
9925a      ............................................................................................--.... 197
NBFu94     ............................................................................................--.... 197
KSS011     ............................................................................................--.... 197
NB0007     ............................................................................................--.... 197
NBP3       ............................................................................................--.... 197
8224       ............................................................................................--.... 197
0028       ..................................................................A...........................--.... 197
0103       ..................................................................A...........................--.... 197
6000       ..................................................................A...........................--.... 197
8641       ..................................................................A...........................--.... 197
NGY122     ....................A.........................................................................--.... 197
2018       ............................................................................................--.... 197
9433       ............................................................................................--.... 197
KJA019     ............................................................................................--.... 197
8656       ............................................................................................--.... 197
9526       ............................................................................................--.... 197
KJA024     ............................................................................................--.... 197
KJA018     ............................................................................................--.... 197
9353       ............................................................................................--.... 197
2009       ............................A.....................................G...........................--.... 197
9648-2     ............................A.....................................G...........................--.... 197
KSS014     ............................A.....................................G...........................--.... 197
KSS015     ............................A.....................................G...........................--.... 197
9593       ............................A.....................................G...........................--.... 197
KSS012     ............................A.....................................G...........................--.... 197
7339       ............................................................................................--.... 197
6746       .......................................................T......................................--.... 197
9371       ............................................................................................--.... 197
EL031      ............................................................................................--.... 197
NGY140     ..........A...C........................................T......................................--.... 198
NGY142     ..........A...C........................................T......................................--.... 198
NGY128-2   ..........A...C........................................T......................................--.... 198
8473       ..........A...................................................................................--.... 198
8539       ..........A...................................................................................--.... 198
BOG0007    ..........A...................................................................................--.... 198
BOG0001    ...A......A............................................T...T..................................--.... 198
9020       ---------------------------------------------..........-.A.................A..G...............CA.... 146
9024       ---------------------------------------------..........-.A.................A..G...............CA.... 146
```

```
                          220                 240                 260                 280                 300
                            |                   |                   |                   |                   |
0098      ACCTCCCGGCGCGGCACGCGCCAAGGAAAACGAACGAGGGCGCGCGCGCGGGCGCCTCCCCCC--GGGGAGGAGGCCGGCGCGCGCCCGCCGAAAACACG 295
NGY128-1  ...............................................................--................................... 295
7006      ...............................................................--................................... 295
8708      ...............................................................--................................... 295
9246      ...............................................................--................................... 295
9312      ...............................................................--................................... 295
9315      ...............................................................--................................... 295
9318      ...............................................................--................................... 295
9524      ...............................................................--................................... 295
9576      ...............................................................--................................... 295
9599      ...............................................................--................................... 295
9601      ...............................................................--................................... 295
9648-1    ...............................................................--................................... 295
9655      ...............................................................--................................... 295
9661      ...............................................................--................................... 295
9668      ...............................................................--................................... 295
9669      ...............................................................--................................... 295
9905      ...............................................................--................................... 295
9925a     ...............................................................--................................... 295
NBFu94    ...............................................................--................................... 295
KSS011    ...............................................................--................................... 295
NB0007    ...............................................................--................................... 295
NBP3      ...............................................................--................................... 295
8224      ...............................................................--................................... 295
0028      ...............................................................C-................................... 296
0103      ...............................................................C-................................... 296
6000      ...............................................................C-................................... 296
8641      ...............................................................C-................................... 296
NGY122    ...............................................................--................................... 295
2018      ...............................................................--................................... 295
9433      ...............................................................--................................... 295
KJA019    ...............................................................--................................... 295
8656      ...............................................................--................................... 295
9526      ...............................................................--................................... 295
KJA024    ...............................................................--................................... 295
KJA018    ...............................................................--................................... 295
9353      ...............................................................--................................... 295
2009      ...................................................C...........--................................... 295
9648-2    ...................................................C...........--................................... 295
KSS014    ...................................................C...........--................................... 295
KSS015    ...................................................C...........--................................... 295
9593      ...................................................C...........--................................... 295
KSS012    ...................................................C...........--................................... 295
7339      ...............................................................C-.................................T. 296
6746      ...............................................................CC.................................T. 297
9371      ...............................................................C-.................................T. 296
EL031     ...............................................................-A.................................T. 296
NGY140    ................T..............................................--.............................-..... 295
NGY142    ................T..............................................--.............................-..... 295
NGY128-2  ................T..............................................--.............................-..... 295
8473      ................A..............................................--.............................T..G.. 296
8539      ................A..............................................--.............................T..G.. 296
BOG0007   ................A..............................................--.............................T..G.. 296
BOG0001   ................T..............................................--.............................T..... 296
9020      ......................................................G........--......CG.....................CGA... 244
9024      ......................................................G........--......CG.....................CGA... 244
```

```
                 320                 340                 360                 380                 400
                   |                   |                   |                   |                   |
0098       CGAGTCAAAACGACTCCCGGCAACGGATATCTCGGCTCTCGCATCGATGAAGAACGTAGCGAAATGCGATACGTGGTGTGAATTGCAGAATCCCGCGAAC 395
NGY128-1   .................................................................................................... 395
7006       .................................................................................................... 395
8708       .................................................................................................... 395
9246       .................................................................................................... 395
9312       .................................................................................................... 395
9315       .................................................................................................... 395
9318       .................................................................................................... 395
9524       .................................................................................................... 395
9576       .................................................................................................... 395
9599       .................................................................................................... 395
9601       .................................................................................................... 395
9648-1     .................................................................................................... 395
9655       .................................................................................................... 395
9661       .................................................................................................... 395
9668       .................................................................................................... 395
9669       .................................................................................................... 395
9905       .................................................................................................... 395
9925a      .................................................................................................... 395
NBFu94     .................................................................................................... 395
KSS011     .................................................................................................... 395
NB0007     .................................................................................................... 395
NBP3       .................................................................................................... 395
8224       .................................................................................................... 395
0028       .................................................................................................... 396
0103       .................................................................................................... 396
6000       .................................................................................................... 396
8641       .................................................................................................... 396
NGY122     .................................................................................................... 395
2018       .................................................................................................... 395
9433       .................................................................................................... 395
KJA019     .................................................................................................... 395
8656       .................................................................................................... 395
9526       .................................................................................................... 395
KJA024     .................................................................................................... 395
KJA018     .................................................................................................... 395
9353       .................................................................................................... 395
2009       .................................................................................................... 395
9648-2     .................................................................................................... 395
7339       .................................................................................................... 396
6746       .................................................................................................... 397
9371       .................................................................................................... 396
EL031      .................................................................................................... 396
9020       ......G............................................................................................. 344
9024       ......G............................................................................................. 344
```

```
                     420                 440                 460                 480                 500
                       |                   |                   |                   |                   |
0098       CATCGAATCTTTGAACGCAAGTTGCGCCCGAGGCCATCCGGCCGAGGGCACGCCTGCCTGGGCGTCACGCCCCCGGTCGCTCCGCGCCGCCCCGTCCCCC 495
NGY128-1   .................................................................................................... 495
7006       .................................................................................................... 495
8708       .................................................................................................... 495
9246       .................................................................................................... 495
9312       .................................................................................................... 495
9315       .................................................................................................... 495
9318       .................................................................................................... 495
9524       .................................................................................................... 495
9576       .................................................................................................... 495
9599       .................................................................................................... 495
9601       .................................................................................................... 495
9648-1     .................................................................................................... 495
9655       .................................................................................................... 495
9661       .................................................................................................... 495
9668       .................................................................................................... 495
9669       .................................................................................................... 495
9905       .................................................................................................... 495
9925a      .................................................................................................... 495
NBFu94     .................................................................................................... 495
KSS011     .................................................................................................... 495
NB0007     .................................................................................................... 495
NBP3       .................................................................................................... 495
8224       .................................................................................................... 495
0028       .................................................................................................... 496
0103       .................................................................................................... 496
6000       .................................................................................................... 496
8641       .................................................................................................... 496
NGY122     .................................................................................................... 495
2018       .................................................................................................... 495
9433       .................................................................................................... 495
KJA019     .................................................................................................... 495
8656       .................................................................................................... 495
9526       .................................................................................................... 495
KJA024     .......................................................................................A............ 495
KJA018     .................................................................................................... 495
9353       .................................................................................................... 495
2009       .......................................................................................A............ 495
9648-2     .......................................................................................A............ 495
KSS014     .......................................................................................A............ 495
KSS015     .......................................................................................A............ 495
9593       .......................................................................................A............ 495
KSS012     .......................................................................................A............ 495
7339       .......................................................................................A............ 496
6746       .......................................................................................A.........T.. 497
9371       .......................................................................................A............ 496
EL031      .......................................................................................A............ 496
NGY140     .......................................................................................A...T........ 495
NGY142     .......................................................................................A...T........ 495
NGY128-2   .......................................................................................A............ 495
8473       .......................................................................................A............ 496
8539       .......................................................................................A............ 496
BOG0007    .......................................................................................A............ 496
BOG0001    .......................................................................................A............ 496
9020       .......................................................................................C.....C..G... 444
9024       .......................................................................................C.....C..G... 444
```

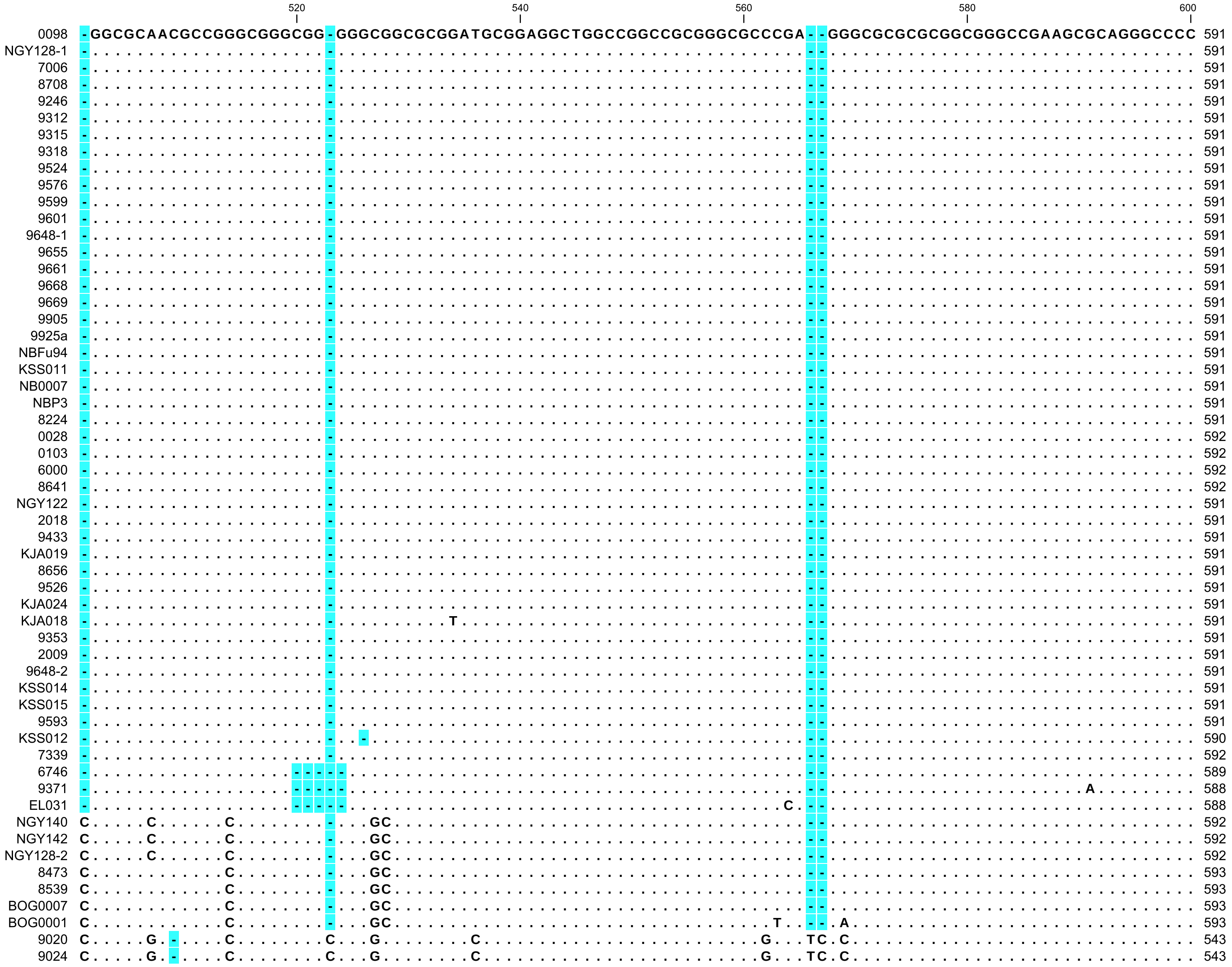

```
                 620                 640                 660                 680                 700
                 |                   |                   |                   |                   |
0098      T-CCGCGGGGCGCGCCGCGGCGAGTGGTGGGATGTCATCGTCGCCGAGGCCGTCGCGCGGGAAGGGGAGAGAGGAACCCCGACGCGAGACCGCGTGCGCT 690
NGY128-1  .-.................................................................................................. 690
7006      .-.................................................................................................. 690
8708      .-.................................................................................................. 690
9246      .-.................................................................................................. 690
9312      .-.................................................................................................. 690
9315      .-.................................................................................................. 690
9318      .-.................................................................................................. 690
9524      .-.................................................................................................. 690
9576      .-.................................................................................................. 690
9599      .-.................................................................................................. 690
9601      .-.................................................................................................. 690
9648-1    .-.................................................................................................. 690
9655      .-.................................................................................................. 690
9661      .-.................................................................................................. 690
9668      .-.................................................................................................. 690
9669      .-.................................................................................................. 690
9905      .-.................................................................................................. 690
9925a     .-.................................................................................................. 690
NBFu94    .-.................................................................................................. 690
KSS011    .-.................................................................................................. 690
NB0007    .-.................................................................................................. 690
NBP3      .-.................................................................................................. 690
8224      .-.................................................................................................. 690
0028      .-.................................................................................................. 691
0103      .-.................................................................................................. 691
6000      .-.................................................................................................. 691
8641      .-.................................................................................................. 691
NGY122    .-.................................................................................................. 690
2018      .-.................................................................................................. 690
9433      .-.................................................................................................. 690
KJA019    .-.................................................................................................. 690
8656      .-.................................................................................................. 690
9526      .-.................................................................................................. 690
KJA024    .-.................................................................................................. 690
KJA018    .-.................................................................................................. 690
9353      .-............................................................................................GT.... 690
2009      .-.................................................................................................. 690
9648-2    .-.................................................................................................. 690
KSS014    .-.................................................................................................. 690
KSS015    .-.................................................................................................. 690
9593      .-.................................................................................................. 690
KSS012    .-.................................................................................................. 689
7339      .-.................................................................................................. 691
6746      .-.................................................................................................. 688
9371      .-.................................................................................................. 687
EL031     .-..............................................................................................T... 687
NGY140    .T.............................T...........T..........C.......C...................A...........C.T... 692
NGY142    .T.............................T...........T..........C.......C...................A...........C.T... 692
NGY128-2  .T.............................T...........T..........C.......C...................A...........C.T... 692
8473      .T.........................................T..................C.............A.....G...........C.T... 693
8539      .T.........................................T..................C.............A.....G...........C.T... 693
BOG0007   .T.........................................T..................C.............A.....G...........C.T... 693
BOG0001   .T.........................................T..................C.............A.....G.............T... 693
9020      .-.........................................T.................GC................G..A.....GA....C.T... 642
9024      .-.........................................T.................GC................G..A.....GA....C.T... 642
```

```
                                     720
                                      |
0098       CGCACCGCGACCCCAGGTCAGGCGGGACTAC 721
NGY128-1   ............................... 721
7006       ............................... 721
8708       ............................... 721
9246       ............................... 721
9312       ............................... 721
9315       ............................... 721
9318       ............................... 721
9524       ............................... 721
9576       ............................... 721
9599       ............................... 721
9601       ............................... 721
9648-1     ............................... 721
9655       ............................... 721
9661       ............................... 721
9668       ............................... 721
9669       ............................... 721
9905       ............................... 721
9925a      ............................... 721
NBFu94     ............................... 721
KSS011     ............................... 721
NB0007     ............................... 721
NBP3       ............................... 721
8224       ............................... 721
0028       ............................... 722
0103       ............................... 722
6000       ............................... 722
8641       ............................... 722
NGY122     ............................... 721
2018       ............................... 721
9433       ............................... 721
KJA019     ............................... 721
8656       ......................T........ 721
9526       .......................A....... 721
KJA024     ............................... 721
KJA018     ............................... 721
9353       ............................... 721
2009       ............................... 721
9648-2     ............................... 721
KSS014     ............................... 721
KSS015     ............................... 721
9593       ............................... 721
KSS012     ............................... 720
7339       ............................... 722
6746       ............................... 719
9371       ............................... 718
EL031      ......................T........ 718
NGY140     ............................... 723
NGY142     ............................... 723
NGY128-2   ............................... 723
8473       ............................... 724
8539       ............................... 724
BOG0007    .......................A....... 724
BOG0001    ............................... 724
9020       ............................C.. 673
9024       ............................C.. 673
```
