## Supporting Information for "Genome diversity and phylogeny of the section *Alatae* of genus *Lemna* (Lemnaceae), comprising the presumed species *Lemna aequinoctialis, Le. perpusilla* and *Le. aoukikusa*"

***New Phytologist* Supporting Information**

The following Supporting Information is available for this article:

**Table S1** The list of duckweed accessions and research conducted on them

**Table S2** List of primers used in this study.

| Primer ID | Sequence, 5'→3' | Direction | Target | Used for |
|---|---|---|---|---|
| DW18S-F1 | CCGCCCGCGACGTCGCGA | Forward | 18S rDNA | ITS1-5.8S rDNA-ITS2 amplification |
| DW25S-R1 | TTATATGCTTAAACTCGGCG | Reverse | 25S rDNA | ITS1-5.8S rDNA-ITS2 amplification |
| TBP-Fex1 | AACTGGGCBAARGGNCAYTAYAC | Forward | β-tub exon1 | β-tub gene intron 1 amplification |
| TBP-Rex1 | ACCATRCAYTCRTCDGCRTTYTC | Reverse | β-tub exon2 | β-tub gene intron 1 amplification |
| TBP-Fin | GARAAYGCHGAYGARTGYATG | Forward | β-tub exon2 | β-tub gene intron 2 amplification |
| TBP-Rin | CRAAVCCBACCATGAARAARTG | Reverse | β-tub exon3 | β-tub gene intron 2 amplification |

**Table S3** Schematic representation of the numerical output of TBP profiles for each analyzed accession

**Fig. S1** Alignment of the *atpF-atpH* spacers of *Le. aequinoctialis, Le. aoukikusa, Le. perpusilla* and *Le. tenera* clones. Matching residues are shown as dots. Gaps are highlighted in light-blue.

**Fig. S2** Alignment of *psbK–psbI* spacers of *Le. aequinoctialis*, *Le. aoukikusa*, *Le. perpusilla* and *Le. tenera* clones. Matching residues are shown as dots. Gaps are highlighted in light-blue.

**Fig. S3** Alignment of ITS1-5.8S-ITS2 sequences of *Le. aequinoctialis*, *Le. aoukikusa*, *Le. perpusilla* and *Le. tenera* clones. Matching residues are shown as dots. Gaps are highlighted in light-blue.
